# High-resolution insights into the geographic differentiation and hybridisation of *Odontobutis* gobies through integrated eDNA and SNP analyses

**DOI:** 10.64898/2025.12.05.692706

**Authors:** Satsuki Tsuji, Yugo Miuchi, Naoki Shibata, Katsutoshi Watanabe

## Abstract

Understanding fine-scale population genetic structure is essential for biodiversity conservation and evolutionary research, but conventional phylogeographic studies often face labour and financial cost constraints. This study proposes a two-step survey strategy that integrates environmental DNA (eDNA) analysis and PCR-based genome-wide SNP genotyping, aiming to evaluate its effectiveness by comprehensively characterising the population structure of widely distributed species. As a model system, we selected the odontobutid gobies, *Odontobutis obscurus* and *O. hikimius*, which occur in western Japan. Initially, water samples were collected from 335 sites across western Japan. Subsequently, tissue sampling for SNP analysis was conducted at 49 sites representing the regional groups and species. The eDNA analysis revealed two major mitochondrial clades within *O. obscurus*, each comprising multiple geographically distinct groups. Subsequently, tissue sampling and SNP analysis were conducted at representative sites of each regional group and species. Nuclear genomic SNP data (661 loci) corroborated the deep divergence between the two clades of *O. obscurus* and, unexpectedly, indicated that *O. hikimius*, whose range lies at their boundary, originated through hybridisation between them. Geographic patterns of the regional groups inferred from both mitochondrial and nuclear data were largely explained by historical geological events such as mountain uplift and ancient river system dynamics, and provide unprecedentedly detailed insight into the population structuring of the focal species. This study demonstrates that the integration of eDNA and SNP analyses provides a cost-effective and scalable approach for high-resolution phylogeographic surveys.

## 1 Introduction

Phylogeography examines the spatial patterns of genetic lineages within a species or related species to infer their distribution mechanisms, providing valuable insights into the evolutionary history of organisms and the processes shaping biodiversity (Avise, 2000; Bermingham and Moritz, 1998; Moritz, 2002; Sersics et al., 2011). Over the past nearly four decades, numerous phylogeographic studies, albeit only a tiny fraction relative to the vastness of biodiversity, have advanced our understanding of species and regional faunal histories (Beheregaray, 2008; Hickerson et al., 2010). Analysing regional population differentiation has also often revealed cryptic species that were morphologically overlooked, significantly contributing to the reassessment of species diversity (e.g. Jense et al., 2024; Knowlton, 1993; Unmack et al., 2023). Nevertheless, expanding human activities continue to cause the loss of such regional genetic groups and species worldwide through habitat destruction, overexploitation and the introduction of invasive species (Almeida-Rocha et al., 2020; Seebens et al., 2018). We may now be facing the critical opportunity to directly investigate the historical trajectories of regional faunas through phylogeographic studies. Thus, there is an urgent global need to accelerate phylogeographic research to comprehensively identify and characterise regional biodiversity elements within species and species groups (Beheregaray, 2008; Turchetto-Zolet et al., 2013). The accumulation of phylogeographic information provides a scientific basis for biodiversity conservation, raising societal awareness and fostering conservation actions (Bowen, 2016; Scoble and Lowe, 2010).

Obtaining reliable phylogeographic data requires sampling numerous individuals from many locations across the species’ range and ideally analysing multiple genome-wide loci (Garrick et al., 2015; Kalkauskas et al., 2021; Liu et al., 2022). Therefore, developing a cost- and labour-efficient survey strategy is key to facilitate phylogeographic research (Pelletier et al., 2022). Recently, biodiversity monitoring using environmental DNA (eDNA), released into the environment by macroorganisms, has been applied to phylogeographic studies (Antich et al., 2023; Pedersen et al., 2021; Tsuji et al., 2023; Turon et al., 2020; Yatsuyanagi et al., 2024). It is expected to become a new survey method due to its simplicity of sampling and analysis processes. Previous research has shown that eDNA-based survey enables large-scale, multi-site sampling over short periods with small teams, simultaneous analysis of multiple species, and the reuse of frozen samples (Tsuji et al., 2025, 2023). However, a current limitation is that the analysable genetic regions are primarily restricted to short mitochondrial DNA (mtDNA) sequences (typically less than 450 base pairs) (Couton et al., 2023; Nakajima and Tsuri, 2024). To obtain detailed population genetic information, it is necessary to analyse numerous genomic loci rather than focusing solely on a single gene (Ballard and Whitlock, 2004). In the last decade, genome-wide single-nucleotide polymorphism (SNP) analysis has increasingly become a common approach in tissue-based phylogeographic studies, e.g., by using reduced-representation genome sequencing methods such as the restriction site-associated DNA sequencing (RAD-seq; Baird et al., 2008) and genotyping by random amplicon sequencing-direct (GRAS-Di; Enoki and Takeuchi, 2018). Several simpler, more cost-effective, but small-scale PCR-based methods have also been developed and are now widely applied [e.g., multiple arbitrary amplicon sequencing (MAAS), Fujimoto et al., 2022; multiplexed ISSR genotyping by sequencing (MIG-seq), Suyama and Matsuki, 2015]. Even with these simpler methods, hundreds to thousands of SNPs can be efficiently detected from small amounts of DNA material. However, these methods still require labour- and time-intensive field surveys for tissue sampling and have a high risk of overlooking locally distributed populations due to the difficulty of dense sampling.

Here, we propose a novel survey strategy that integrates eDNA analysis and PCR-based genome-wide SNP analysis, aiming to maximise the strengths of each method while addressing their respective limitations. This approach has the potential to enable more efficient and comprehensive phylogeographic surveys that are difficult to achieve with conventional approaches. In this survey strategy, eDNA analysis is initially employed to conduct labour- and cost-efficient, high-density surveys across the entire distribution range of the target species, thereby identifying regional genetic groups and providing an overview of their distribution patterns. Subsequently, representative sites are selected to reflect the distribution of the identified regional groups, followed by capture surveys and genome-wide SNP analysis. This two-step approach is expected to minimise the risk of overlooking locally distributed groups or redundant sampling of populations from the same groups, while also reducing analytical effort and costs. Moreover, since eDNA concentration is positively correlated with target biomass (Rourke et al., 2022), it may help identify sites where capture surveys can be conducted more effectively.

We selected odontobutid gobies, a group widely distributed across western Japan, as a suitable model for demonstrating the effectiveness of our novel phylogeographic survey strategy proposed herein. The Japanese archipelago, an island system extending over 2000 km in a northeast–southwest direction along the eastern margin of East Asia, harbours a unique and heterogeneous biota that has likely been shaped by historical geographic and climatic changes since its formation in the Neogene (Nakagawa et al., 2016; Watanabe, 2012; Yonekura, 2001). It has long served as a valuable setting for phylogeographic studies (e.g. Motokawa and Kajihara, 2017). In particular, freshwater organisms, due to their low dispersal ability, exhibit various patterns of geographic differentiation within species and closely related species groups, providing excellent opportunities for phylogeographic studies to explore the relationship between organismal distributions and the tectonic history of the archipelago. In Japan, two freshwater species of the odontobutid gobies are recognised: *Odontobutis obscurus* (Temminck & Schlegel, 1845), which is widely distributed across western Japan, and *Odontobutis hikimius* Iwata and Sakai, 2002, which is confined to a limited area within the range of the former. Previous studies based on allozyme and mtDNA analyses have reported that *O. obscurus* includes at least five phylogeographic groups (Mukai and Nishida, 2003; Sakai et al., 1998). *Odontobutis hikimius* was once considered a regional group of *O. obscurus*, but is now recognised as a morphologically and genetically distinct species, which is locally distributed in the upper reaches of the Takatsu River system and adjacent river basins (Fig. 1; Iwata and Sakai, 2002). However, due to limited geographic sampling and the low resolution of previously used genetic markers, the overall phylogeographic patterns of these two species, including their evolutionary relationships, remain insufficiently resolved. This highlights the need for further surveys involving denser sampling and genome-wide genetic analyses across their entire distribution ranges.

**Fig. 1.**
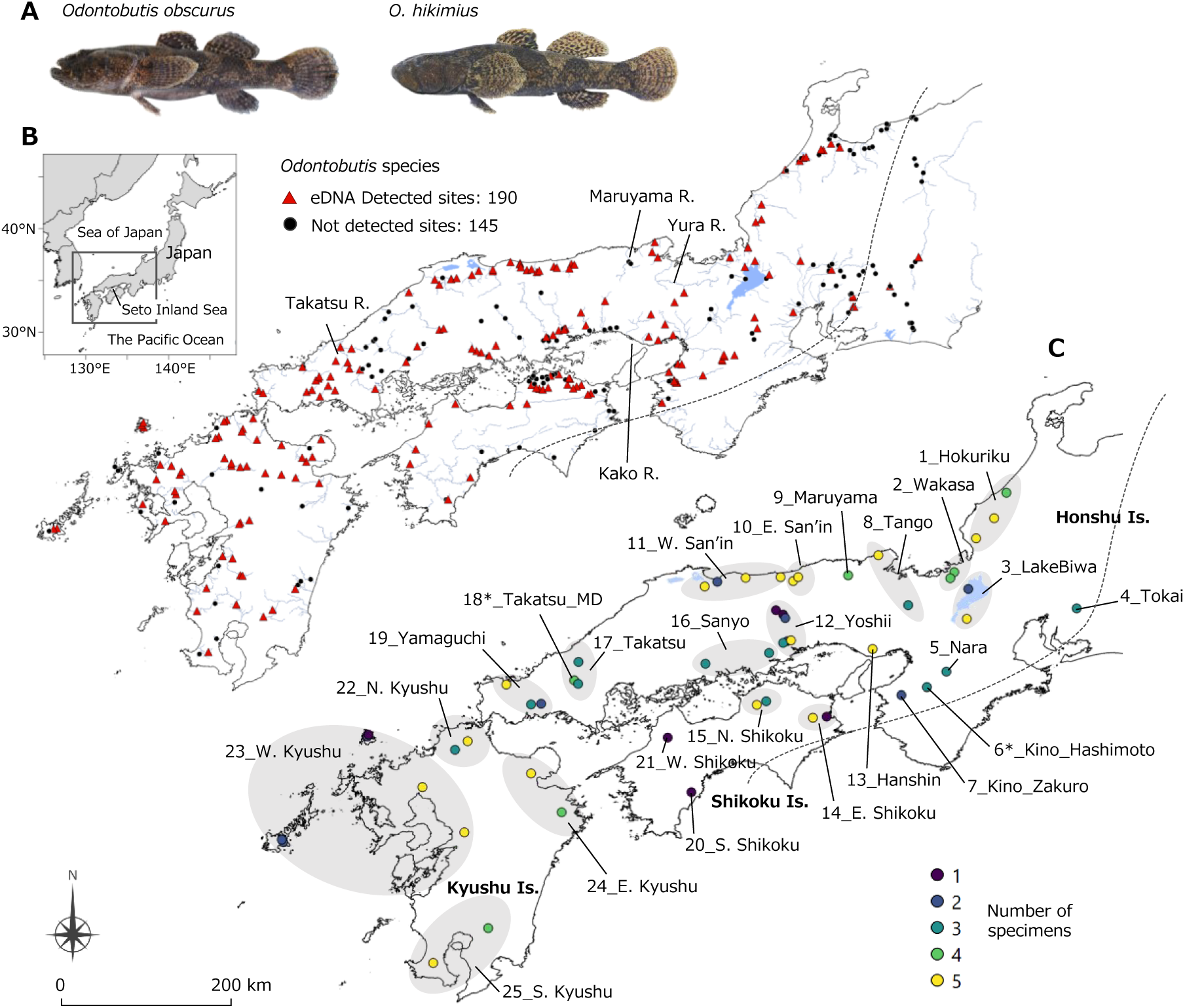
The target *Odontobutis* species and geographic distribution of sampling sites. (A) *O. obscurus* (left) and *O. hikimius* (right). (B) Water sampling sites for eDNA analysis. Red triangles and black circles indicate the detection and non-detection sites of *Odontobutis* eDNA, respectively. The dashed black line indicates the boundary of the natural distribution range of *Odontobutis* species. (C) Capture survey sites for tissue sampling. Coloured circles indicate the number of specimens used for tissue DNA analysis. The map scale shared between panels (B) and (C)

This study aims to demonstrate the effectiveness of the novel survey strategy that combines eDNA and PCR-based genome-wide SNP analyses, by applying it to comprehensively elucidate the population structure of *O. obscurus* and *O. hikimius* in Japan. In the first step, eDNA analysis targeting mtDNA was conducted to fully identify regional genetic groups of the two species and to estimate their distributions across the species’ entire ranges. In the second step, based on the distribution patterns of these regional groups, we selected sampling sites and conducted SNP and extended mtDNA analyses using tissue-derived DNA from specimens collected at these sites. These analyses successfully reconstructed high-resolution population structures of the two species, revealing two major genetically distinct groups with partially overlapping distributions in *O. obscurus*. Furthermore, the demographic modelling approach provided insights into the evolutionary relationships between the two species, especially regarding the origin of *O. hikimius*, which is locally distributed around the boundary area of the two *O. obscurus* groups.

## 2 Materials and methods

### 2.1 Target species

*Odontobutis obscurus* and *O. hikimius* are freshwater goby species distributed in Japan (Hosoya, 2019; Iwata and Sakai, 2002). *Odontobutis obscurus* is also found on Geoje Island, South Korea, where it is now nearly extinct (Hosoya, 2019). The common ancestor of these species is inferred to have diverged from continental congeners in the Late Miocene, concurrent with the formation of the Japanese archipelago, and to have later split into the two species in the Pliocene (Hu et al., 2023). Both species grow to approximately 15–25 cm in total length and are voracious predators, feeding on crustaceans, fish, and aquatic insects. While *O. hikimius* is locally distributed around a specific water system (Takatsu River system; Fig. 1B), *O. obscurus* is widely distributed throughout western Japan, inhabiting slow-flowing environments such as midstream sections of rivers, irrigation channels, and ponds. Previous studies have identified five genetically distinct regional groups in *O. obscurus* (W. Seto, E. Seto, W. Kyushu, E. Kyushu, and San’in-Biwa-Ise; Sakai et al., 1998), although the additional groups may also exist, as suggested by Mukai and Nishida (2003).

### 2.2 Environmental DNA sample collection

Water sampling for eDNA analysis was carried out at a total of 335 sites across western Japan between 2017 and 2024 (Fig. 1B; Table S1). Of these, 91 samples collected between 2017 and 2021 were used in a previous study for eDNA-based phylogeographic analyses of five species, including the two *Odontobutis* species (Tsuji et al., 2023). In this study, we combined the data from these earlier samples with additional data obtained from surveys newly conducted between 2021 and 2024 to perform further analyses. At all sites, one litre of surface water was collected using a disposable plastic cup and water bag (DP16-TN1000, Yanagi, Aichi, Japan). To prevent the degradation of eDNA, 0.5 mL of benzalkonium chloride (BAC, 10% w/v; Osvan S; Nihon Pharmaceutical Co., Ltd, Tokyo, Japan) was added to the collected water and agitated well (Yamanaka et al., 2017). The water sample was filtered immediately in the car while in transit using a GF/F glass fibre filter (47 mm diameter, 0.7 μm mesh size; GE Healthcare Japan, Tokyo, Japan) and a filter holder. To ensure the reliability of the results, as a field negative control (FNC), one litre of ultrapure water with 0.5 mL of BAC was filtered at the end of each survey day using the same method and the same equipment as for the environmental samples. After filtration, all filter samples were immediately frozen by dry ice or –20°C freezer.

DNA was extracted from each filter sample using a modified spin column-based protocol with the DNeasy Blood and Tissue Kit (Qiagen), following Tsuji et al. (2024). Briefly, filters with DNA were lysed with proteinase K, and DNA was purified using DNeasy columns. The extracted eDNA was eluted in Buffer AE and stored at −20°C until analysis. Full protocol details are provided in the Supplementary file.

### 2.3 Paired-end library preparation for eDNA samples

The eDNA library preparation followed the protocol of Tsuji et al. (2023), targeting a 366 bp fragment of the mtDNA 12S ribosomal RNA (rRNA) using *Odontobutis* group-specific primers with internal standard DNAs. After two rounds of PCR and size selection, indexed libraries were pooled and sequenced on an Illumina NovaSeq 6000 (250 bp paired-end). Detailed procedures and primer information are provided in the Supplementary file.

### 2.4 Bioinformatics processing and denoising for eDNA sequence data

Bioinformatic processing of eDNA sequence data followed the procedure described in Tsuji et al. (2023). Denoising was performed using the Divisive Amplicon Denoising Algorithm 2 (DADA2) package (v. 1.22; Callahan et al., 2016), and amplicon sequence variants (ASVs) were quantified based on internal standard DNA calibration. Species assignment was conducted via BLASTN using a custom reference database. A two-step filtering procedure was applied to minimise false positives (Tsuji et al., 2023), and the resulting ASVs were treated as haplotypes for phylogenetic analyses. Full details of the processing steps are provided in the Supplementary file.

### 2.5 Phylogeographic analyses and visualisation

Phylogeographic analyses of the eDNA data were conducted using maximum likelihood (ML) tree inference and haplotype network estimation. An ML tree was inferred with IQ-TREE v. 2.4.0 (Minh et al., 2020) for all haplotypes that passed the data-denoising steps, the reference sequences (Fig. 2A; Mukai and Nishida, 2003) and outgroups (*Micropercops swinhonis* and *Odontobutis potamophilus*). The optimal substitution model was determined using ModelFinder (Kalyaanamoorthy et al., 2017) and the TN+F+G4 model was selected based on the corrected Akaike Information Criterion (AICc). Node support was assessed using the Ultrafast Bootstrap approach (Hoang et al., 2018) with 1000 replicates. The resulting tree was visualised using interactive Tree of Life (iToL) v. 6 (https://itol.embl.de/). A haplotype network was constructed using POPART v. 1.7 (http://popart.otago.ac.nz; Leigh and Bryant, 2015) with the Templeton– Crandall–Sing (TCS) algorithm (Clement et al., 2000). Genetic groups were identified based on the 2-step clades derived from the network following the nesting rule (Templeton and Sing, 1993), which most effectively captured regional genetic structure at the highest resolution. The geographic distribution of these mtDNA groups was visualised using R v. 4.1.1 (R Core Team, 2023) with the packages maps v. 3.4.0 (Becker et al., 2021), mapdata v. 2.3.0 (Brownrigg, 2018), and mapplots v. 1.5.1 (Gerritsen, 2018).

**Fig. 2.**
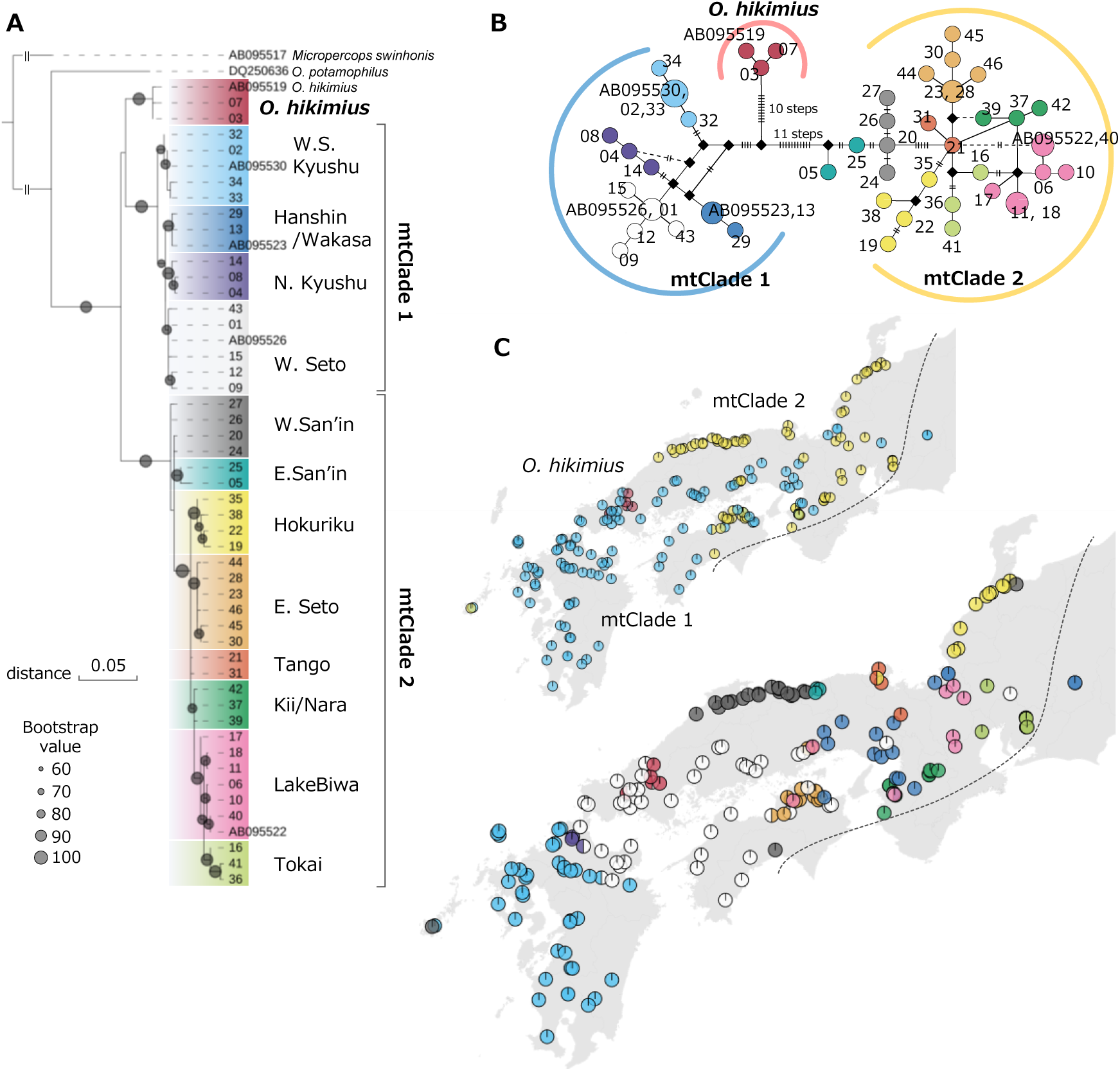
Phylogeny and genetic population structure of Japanese *Odontobutis* species inferred from eDNA analysis of the mtDNA 12S rRNA sequences (366 bp). Regional groups were defined using a two-step nesting of clades (Templeton and Sing, 1993). The colour scheme used for each group is consistent across all panels. (A) Maximum likelihood tree constructed from detected haplotypes (Haplotype ID; Table S5), with reference haplotypes (accession No. in NCBI). Bootstrap support values (> 60%) are indicated by circle sizes. (B) Haplotype network including both detected and reference haplotypes. Short bars indicate the number of base differences between neighbouring haplotypes. Black diamonds indicate missing haplotypes. (C) Distribution map of each regional group. Pie charts represent the relative frequencies of detected groups. The black dashed line indicates the boundary of the natural distribution range *Odontobutis* species

### 2.6 Tissue DNA sample collection

Tissue samples were collected from 49 sites (Table S2, Fig. 1C) and used for sequencing a longer mtDNA region and for nuclear SNP analyses using the MAAS method. The longer mtDNA sequences were employed to improve the reliability of the phylogenetic tree constructed from eDNA data and to estimate divergence times. These collection sites were selected to include at least two or more sites for each of the mtDNA groups identified through the eDNA analysis. For the groups with relatively wide distribution ranges, geographically distant sites were preferentially selected. For an outgroup in MAAS analysis, an individual of *O. potamophilus* from a non-native population in the Tone River, Japan, was added to the analysis. At each site, 1–5 individuals were captured (177 individuals in total; 163 *O. obscurus* + 6 *O. hikimius* + 7 suspected hybrids + 1 *O. potamophilus*; Table S2). Fin clips from each individual were preserved in 99% ethanol and subsequently used for DNA extraction. Tissue DNA was extracted using spin columns (EconoSpin, EP-31201; GeneDesign, Inc., Japan) and reagents provided in the DNeasy Blood and Tissue Kit (Qiagen).

### 2.7 Longer mtDNA sequencing and divergence time estimation

To validate the presence of the mtDNA groups detected by the eDNA analysis and to estimate divergence times among lineages, a longer mtDNA region (950–955 bp; 12S rRNA–tRNA-Val– 16S rRNA), which includes the region targeted in eDNA analysis, was sequenced for a total of 176 specimens from 48 sites (163 *O. obscurus* + 6 *O. hikimius* + 7 suspected hybrids) using either high-throughput amplicon sequencing or Sanger sequencing (Table S2). Detailed procedures and primer information are provided in the Supplementary file.

To estimate the divergence time among regional mtDNA groups, a time-calibrated phylogenetic tree was inferred based on aligned 961 bp sequences obtained from tissue DNA using BEAST v. 2.6.7 (Bouckaert et al., 2019). A total of 83 haplotypes were identified using DnaSP v. 6.12.03 (Rozas et al., 2017) from sequences successfully obtained from 169 specimens across 46 sites (163 *O. obscurus* + 6 *O. hikimius*). Among them, 34 representative haplotypes were selected to encompass all mtDNA regional groups, to reduce computational load and improve parameter convergence in the BEAST analysis (Table S2). As the outgroup, sequence data of three continental congeners (*Odontobutis sinensis*, MZ265331; *O. potamophilus*, MK408452; *O. yaluensis*, KM277942) were obtained from NCBI and included in the analysis. Sequences were aligned using MAFFT (Katoh et al., 2019). To ensure the use of only reliable sites for divergence time estimation, alignment positions with reliability scores below 0.9, as calculated by GUIDANCE2 (Penn et al., 2010), were masked with “N”. The substitution and site models were determined from the six basic models (JC, K80, HKY, TN, SYM and GTR) using ModelFinder based on AICc. The GTR + F + I + G4 model was selected as the best-fit model. The clock model was set to a Relaxed Clock Log Normal model, and the tree prior was set to the Birth–Death model. Three calibration points were employed based on the studies by Hu et al. (2023): (1) the origin time of *Odontobutis,* defined as the period between the most recent common ancestor (MRCA) of all *Odontobutis* species and the divergence time from its sister genus *Neoodontobutis* [log normal distribution, mean = 9.0 million years (Ma), stdev = 5.79; “use originate” setting applied]; (2) the MRCA of *O. sinensis*, *O. potamophila* and *O. yaluensis* (normal distribution, mean = 8.0, stdev = 2.0); and (3) the MRCA of *O. potamophila* and *O. yaluensis* (normal distribution, mean = 3.8, stdev = 1.2). Species outside *Odontobutis* (e.g. *Neoodontobutis*) were not included as outgroups because their inclusion increased alignment ambiguity and destabilised the tree topology. Markov chain Monte Carlo (MCMC) sampling was run for 50 million generations with parameters logged every 5,000 generations. Parameter convergence was assessed using Tracer v. 1.7.2 (Rambaut et al., 2018), ensuring all effective sample size (ESS) values exceeded 200. The initial 10% of trees were discarded as burn-in, and the remaining trees were summarised into a single tree using TreeAnnotator.

### 2.8 MAAS library construction and sequencing

For nuclear SNP analysis of 177 specimens from 49 sites (163 *O. obscurus* + 6 *O. hikimius* + 7 suspected hybrids + 1 *O. potamophilus*), we employed the MAAS method, one of the PCR-based genome-reduction methods (Fujimoto et al., 2022). A MAAS library was prepared based on the protocol described by Fujimoto et al. (2022), with slight modifications as follows. The 1st PCR was performed using four primer sets with adapter sequences at an annealing temperature of 38°C (Table S3). The Multiplex PCR Assay Kit ver. 2 (Takara, Shiga, Japan) was employed as a total of 10 µL PCR reaction mixture containing 1 µL of template DNA. The 1st PCR products were diluted 50-fold in TE buffer (pH 8.0) and used as templates for the 2nd PCR. The 2nd PCR was conducted using primers that included indexing sequences (Hamady et al., 2008) with adapter sequences. The KAPA HiFi HotStart ReadyMix was employed as a total of 12 µL PCR reaction mixture containing 0.3 µM of each index primer and 1 µL of diluted 1st PCR product. Unique indexes were applied to each sample to distinguish them from one another. The thermal conditions were as follows: 3 min at 95°C, 12 cycles of 20 s at 98°C, 15 s at 60°C and 30 s at 72°C, and 5 min at 72°C. Following amplification, fragments within the size range of 300–700 bp were isolated and purified using Sera-Mag SpeedBeads. Fragment size and final library concentration were verified using the Agilent 2200 TapeStation (Agilent Technologies, Santa Clara, CA) and the Invitrogen Qubit 4 Fluorometer (Thermo Fisher Scientific). The final library was shipped to Novogene Co., Ltd. and sequenced on an Illumina NovaSeq X Plus using their gigabyte data purchase service (150 PE sequences).

### 2.9 MAAS data treatment and SNP detection

The raw sequencing data containing primer sequences were removed using cutadapt v4.9 with the -b and -B options specified. Quality control was performed using fastp v0.24.0 (Chen et al., 2018). Residual adapter sequences, low-quality bases (<Q30), and polyG sequences (>10) were removed, and all reads were standardised to a length of 90 bases. SNP detection was performed using the denovo_map.pl pipeline in Stacks v2.68 (Rochette et al., 2019) in paired-end mode. The parameters were set as follows: the maximum allowable distance between stacks (M) = 5 and the minimum depth of coverage required to form a stack (m) = 5. Output files for downstream analyses were generated using the populations program in Stacks with the following settings: a minimum percentage of variant sites shared by all individuals (R) = 0.7 and output restricted to the first SNP per locus (--write-single-snp). Variants with heterozygosity exceeding 75% (--max-obs-het = 0.75) or fewer than two minor alleles (--min-mac = 2) were excluded. All other parameters were left at their default settings.

### 2.10 Population genetic structure and divergence scenarios based on SNPs

To estimate phylogenetic relationships based on nuclear SNPs, a PHYLIP-format file with 688 SNPs obtained from 170 specimens (163 *O. obscurus* + 6 *O. hikimius* + 1 *O. potamophilus*) was generated from the VCF file produced by the populations program, using the Python script vcf2phylip.py (https://github.com/edgardomortiz/vcf2phylip.git). Subsequently, model selection and phylogenetic tree construction were carried out using ModelFinder and IQ-TREE2 with ascertainment bias corrections (ASC). The K2P+ASC+G4 model was selected as the best model based on AICc.

In addition, to estimate the population structure of *O. obscurus* and *O. hikimius*, principal component analysis (PCA) was conducted using PLINK v1.90 (Purcell et al., 2007). The PHYLIP-format input file with 663 SNPs obtained from 176 specimens (163 *O. obscurus* + 6 *O. hikimius* + 7 suspected hybrids) was generated using the same method as described above. Individual admixture proportions were estimated through unsupervised clustering with ADMIXTURE v1.3.0 (Alexander et al., 2009), employing a likelihood-based model. Analyses were performed for a range of genetic clusters (K) from 1 to 15, with the convergence criterion set at 10 (C = 0.0001). To estimate optimal K based on the lowest mean cross-validation error (CV-error), each analysis was repeated 100 times with different random seed values. Data summarisation and visualisation of CV-error values and admixture proportions were performed using an original script in Python v3.12 and R v4.3.3 (R Core Team, 2023). To infer relationships among regional population samples, an individual-level network was also constructed based on the SNP data using SplitsTree v6.4.11 (Huson and Bryant, 2024). The VCF files obtained from 176 specimens used in the above analysis were converted into the nexus files using the Python tool (vcf2phylip.py) and subsequently employed for network analysis. The Neighbour-net method was applied using uncorrected p-distances.

To estimate divergence scenarios for the major lineages of *O. obscurus* and *O. hikimius*, a coalescent simulation-based population demographic analysis was performed using an approximate Bayesian computation approach (ABC) with supervised machine learning based on random forests. For the analysis, DIYABC-RF v1.1.28-14e1552 and abcranger v1.16.67 were used (Collin et al., 2021). The input files with 458 SNPs obtained from 169 specimens (163 *O. obscurus* + 6 *O. hikimius*) were generated from the VCF file produced by the populations program (with options: −*r* 0.5, -*p* 3), using the Python script vcf2phylip.py. It was assumed that there was no bias in the sex ratio of the specimens. The minor allele filtering was omitted, as it had already been addressed during the stacks processing. Four divergence scenarios were set to represent different phylogenetic relationships and gene flow patterns among Japanese *Odontobutis* species (Fig. 4E). Specifically, these include three possible phylogenetic trees for three ncClades, including *O. hikimius*, and a tree suggesting *O. hikimius* has a hybrid origin from two *O. obscurus* ncClades, as indicated by Neighbour-net and Admixture analyses. The prior distributions of demographic and temporal parameters were assigned considering the divergence time estimated from mtDNA data and the assumed generation time: effective population sizes, log-uniform distribution between 10 and 100,000; divergence time, log-normal distributions, with ranges of 100,000–10,000,000 generations for the first split, 10,000–10,000,000 generations for the second, and 10–5,000,000 generations for the most recent split; the mixing ratio, log-uniform distribution between 0.05 and 0.95. Although the exact generation time of *Odontobutis* species remains unknown, we assumed it to be three to four years, based on the observation that *O. obscurus* reaches sexual maturity within one year in captivity (Mashiko and Yamagishi, 1976) and has an estimated lifespan of five to ten years in the wild (Matsuda, 2024). A total of 90,000 datasets were simulated to train the random forest model. The most likely divergence scenario and associated parameters were estimated by a voting mechanism, using 3,000 simulations for training and generating 500 trees per analysis (Collin et al., 2021).

## 3. Results

### 3.1 Population structure outlined by eDNA analysis

Among the 335 water sampling sites, eDNA of either *Odontobutis obscurus* or *O. hikimius* was detected at 190 sites (Tables S1, S4, Fig. 1B). For *O. obscurus*, two highly differentiated clades (mtClade 1 and mtClade 2) were identified, differing by 20 or more nucleotide substitutions (5.5% in sequence differences; Table S5, Fig. 2). The distribution ranges of mtClade 1 and mtClade 2 were mostly distinct, but showed overlap in northeastern Shikoku Island (Kagawa Prefecture) (Figs. 1C, 2C). mtClade 1 comprised four regional groups, corresponding to those identified in previous studies based on both sequence characteristics and geographic distribution (Fig. 2C; Mukai and Nishida, 2003; Sakai et al., 1998; Tsuji et al., 2023). mtClade 2 corresponded to the San’in/Biwa/Ise group reported in previous studies (Mukai and Nishida, 2003; Sakai et al., 1998) and consisted of eight regional groups, including the newly recognised Tango group. The distribution areas of each regional group were generally well delineated (Fig. 2C). However, some haplotypes belonging to the W. Seto (st. 103, 158, 30), W. San’in (st. 202, 274), Lake Biwa (st. 018, 091, 095), Hokuriku (st. 109) and Hanshin/Wakasa (st. 169, 170; known as introduced; Yagyu et al., 2021) groups were found at geographically distant and isolated sites. The ML tree suggested that mtClade 1 and *O. hikimius* together formed a monophyletic group, but its bootstrap support was low (<1.2%; Fig. 2A).

### 3.2 Divergence times inferred from longer mtDNA sequences

To resolve the phylogenetic relationships among the mtDNA groups and estimate their divergence times, 920-bp sequences of mtDNA spanning the 12S rRNA to the 16S rRNA gene regions were obtained from 169 specimens from 48 sites (Table S2). The time-calibrated tree supported the monophyly of each of the two major clades (mtClade 1 and mtClade 2), and its topology closely resembled that obtained from the eDNA analysis (Fig. 3). The results suggested that *O. obscurus* was not monophyletic: mtClade 1 and *O. hikimius* together formed a monophyletic group (BPP = 0.94), which was sister to mtClade 2. The time to the most recent common ancestor (tMRCA) of *O. obscurus* and *O. hikimius* was estimated at 3.90 Ma [95% highest posterior density (HPD), 2.06–5.90 Ma]. The divergence between mtClade 1 + *O. hikimius* was estimated at 3.13 Ma (1.59–4.76). The tMRCA of mtClade 1 was estimated at 1.23 Ma (0.57–1.93). mtClade 2 was suggested to have diverged into the San’in groups (eastern and western) and other groups at 1.99 Ma (0.99–3.10). The Lake Biwa and Tokai groups in mtClade 2, which are separated by the Ibuki–Suzuka Mountains, were suggested to have diverged at 0.44 Ma (0.19–0.72).

**Fig. 3.**
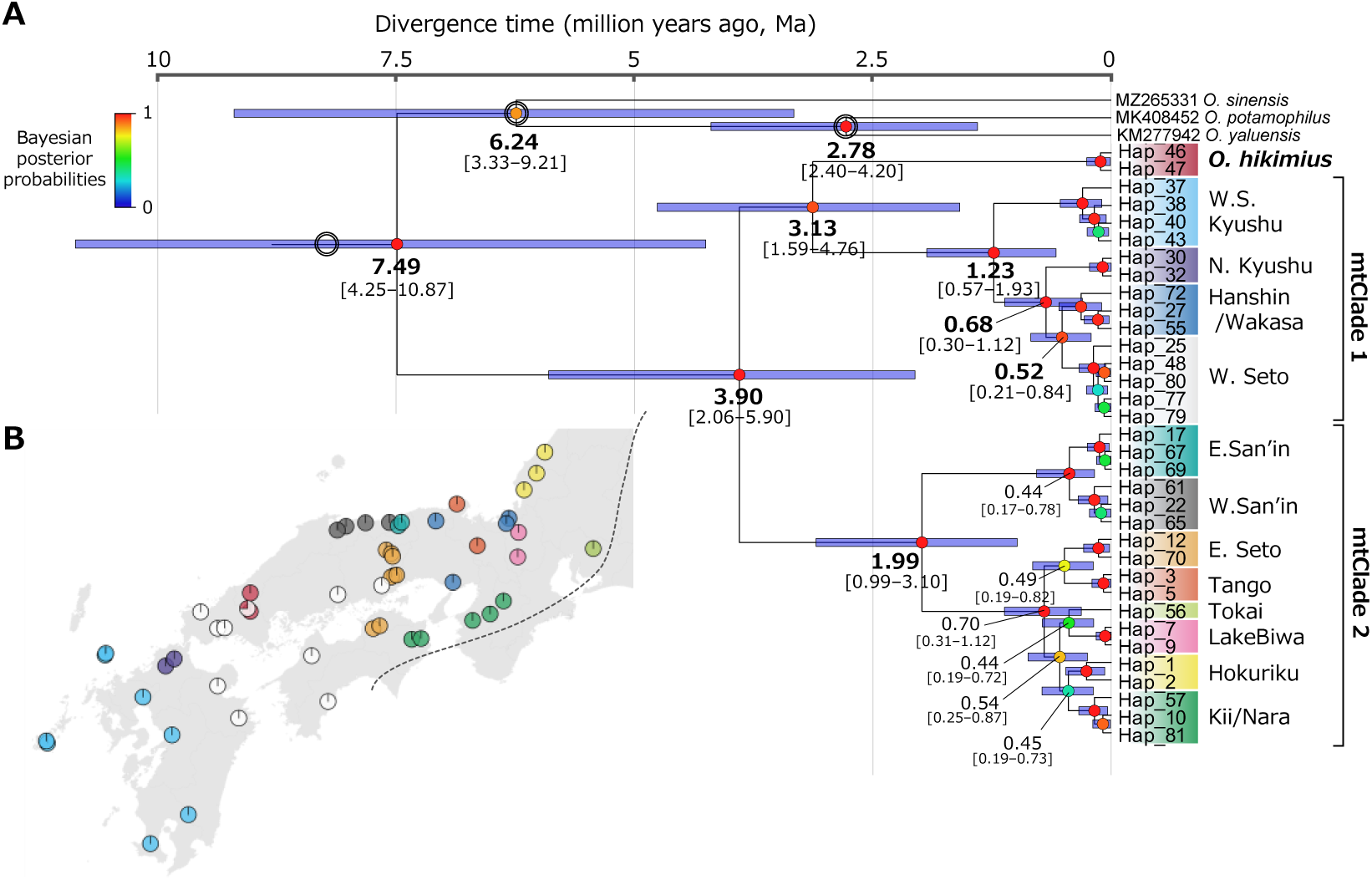
Phylogeny and genetic population structure of Japanese *Odontobutis* species inferred from mtDNA 12S–16S rRNA sequences (aligned 961 bp) obtained from tissue DNA. Colour coding for each regional group corresponds to that in Fig. 1. (A) Time-calibrated Bayesian phylogenetic tree showing estimated divergence times among *Odontobutis* species. The mean estimated divergence time and its 95% highest posterior density (HPD) are shown as numerical values and purple bars. The colour level of the node points indicates Bayesian posterior probabilities (BPPs). Double circles indicate the calibration points based on Hu et al. (2023). (B) Geographic distribution of each regional group based on tissue-derived mtDNA. The pie chart indicates the relative frequency of each haplotype. The black dotted line denotes the natural distributional boundary of *Odontobutis* species.

### 3.3 Population structure inferred from nuclear SNPs

The phylogenetic tree, PCA, and Neighbour-net analyses based on SNP data from *O. obscurus* (n = 163 from 44 sites) and *O. hikimius* (n = 6 from two sites) supported deep genetic differentiation between the two species, as well as the presence of two distinct clades within *O. obscurus* (ncClade 1 and ncClade 2; Table S2, Fig. 4). These clades corresponded to the two clades revealed by the mtDNA analysis based on both eDNA and tissue DNA.

**Fig. 4.**
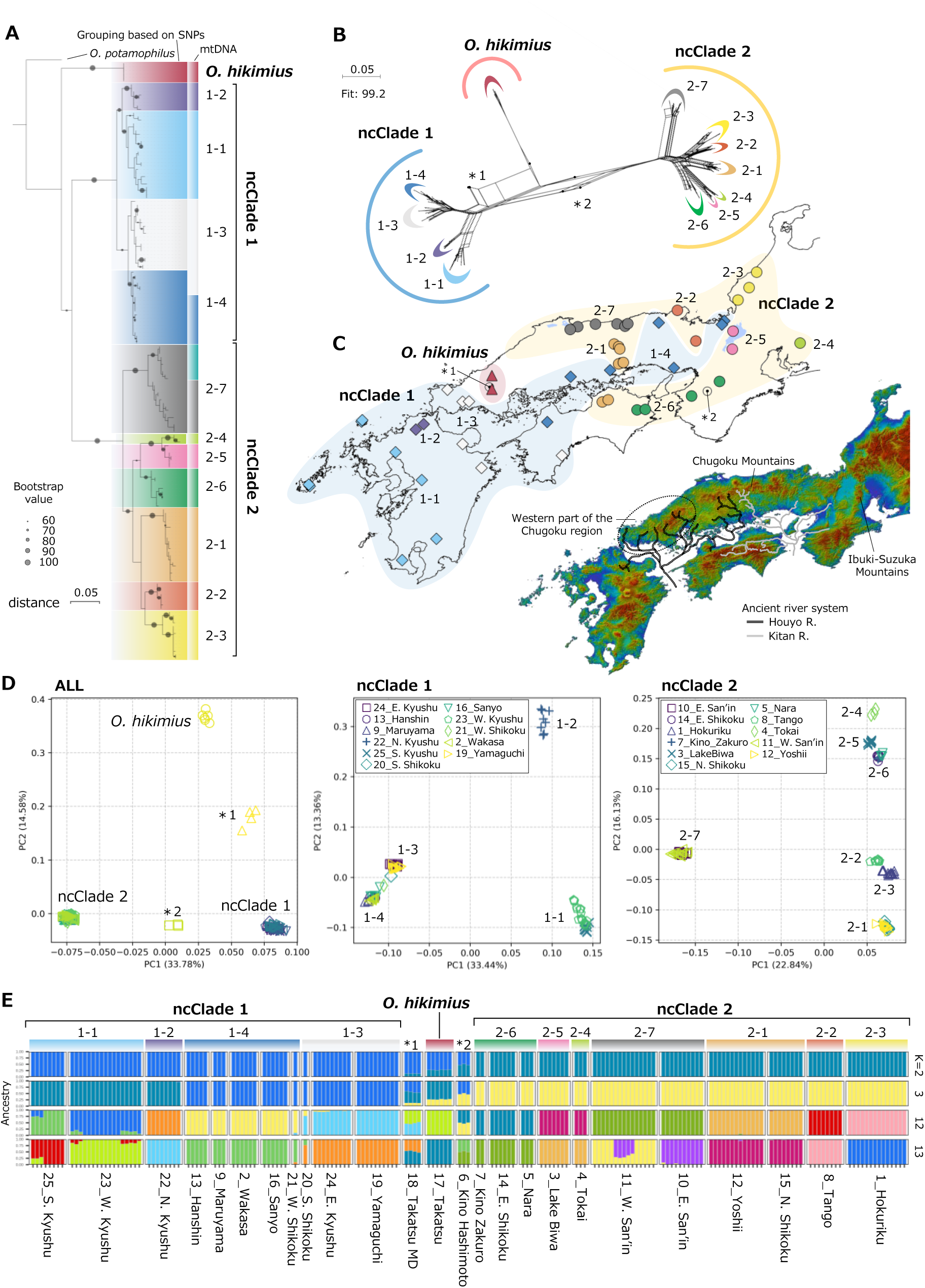
Phylogeny and genetic population structure of Japanese *Odontobutis* species inferred from genome-wide SNPs. (A) Maximum likelihood tree constructed using 688 SNPs. Branch colours indicate groupings based on genome-wide SNPs (left) and mtDNA haplotypes (right). (B) Neighbour-net network based on 663 SNPs. (C) Distribution map of each regional group (upper) and the major geographic barriers and reconstructed flow paths of ancient river systems in the Seto Inland Sea region (lower). (D) Principal component analysis (PCA) based on 663 SNPs: all samples (left), ncClade 1 (centre) and ncClade 2 (right). (E) Ancestry bar plots from ADMIXTURE analysis based on 663 SNPs. The number of clusters was determined based on the number of major lineages (K = 3), SNP-based grouping (K = 12) and eDNA-based grouping (K = 13). Asterisks (*1 and *2) in panels (b)–(d) indicate sampling sites 18_Takatsu_MD and 6_Kino_Hashimoto, respectively (see Fig. 1C). Colour coding of phylogeographic groups is consistent across panels (A) to (C).

The SNP-based phylogenetic tree indicated that the two clades within *O. obscurus* formed a monophyletic group, in contrast to the mtDNA-based tree (Fig. 4A). However, the SNP tree showed an almost trifurcated topology among the two clades and *O. hikimius*, and support for the monophyly of ncClades 1 and 2 was weak (bootstrap value = 69%). The regional groups identified by SNP and mtDNA data are highly concordant. However, the W. San’in and E. San’in groups, which were recognised as distinct regional groups based on mtDNA data, were identified as a single group (ncClade 2-7) in SNP-based analyses (Fig. 4A). In addition, some samples (st. 21_W. Shikoku and st. 16_Sanyo) with mtDNA haplotypes belonging to the W. Seto group of mtClade 1 were placed in ncClade 1-4 rather than in ncClade 1-3, where the other samples were placed (Fig. 4A). PCA revealed several distinct clusters corresponding to the regional groups identified in the SNP-based ML tree (Fig. 4D). These clusters were also supported by the Neighbour-net analysis (Fig. 4B).

The distribution areas of ncClades 1 and 2 were not overlap, but ncClade 1-4 shows a complex geographical distribution pattern at the boundary with the distribution area of ncClade 2. Also for each regional group, the distribution areas were clearly separated. Several groups (ncClade 1-3, 1-4, 2-1 and 2-6) spanned regions that were geographically close but divided by the Seto Inland Sea (Figs. 1B, 4C).

The admixture analysis indicated that increasing the number of clusters resulted in a progressive decrease in the CV-error value, making it difficult to identify an optimal number of clusters (Fig. S1A). Guided by the ML tree, PCA, and Neighbour-net results, K = 3 (corresponding to *O. hikimius* and two clusters for *O. obscurus*), K = 12 (*O. hikimius*, four regional groups of ncClade 1, and seven of ncClade 2), and K = 13 (*O. hikimius* and 12 regional groups for *O. obscurus* as inferred from mtDNA) were considered biologically meaningful clustering (Figs. 4E, S1B). At K = 3, ncClade 1 was divided into two clusters: one comprising ncClades 1-1 and 1-2, and the other comprising ncClades 1-3 and 1-4. Unexpectedly, *O. hikimius* was inferred to consist of genetic contributions from the former cluster and from ncClade 2. A few regional groups (6*_Kino_Hashimoto and 18*_Takatsu_MD) were also found to have genetic contributions from multiple clusters. Separate admixture analyses of ncClade 1 and ncClade 2 confirmed that each clade was clearly divided into regional groups, corresponding to the clusters inferred from the PCA results (four and seven, respectively; Figs. S2, S3).

Whereas *O. hikimius*, unambiguously identified based on both mtDNA and morphological characteristics, was found in the upper reaches of the Takatsu River system (17_Takatsu), four specimens collected from the middle reaches (18*_Takatsu_MD) possessed either *O. hikimius* mtDNA (one specimen) or mtDNA belonging to the W. Seto group (three specimens). SNP-based PCA, Neighbour-net, and admixture analyses indicated that these specimens were genetically intermediate between *O. hikimius* and ncClade 1, suggesting their hybrid status (Table S2, Fig. 4). Also, three specimens from the Hashimoto River, a tributary of the Kino River system (6*_Kino_Hashimoto), had mtDNA of the Kii/Nara group, but all SNP-based analyses indicated that they were genetically intermediate between ncClades 1 and 2 (Table S2, Fig. 4).

### 3.4 Divergence and demographic history

The origin and divergence patterns among the two highly differentiated groups within *O. obscurus* and *O. hikimius* were examined using the DIYABC-RF model selection based on 458 SNPs. Three trifurcating divergence scenarios (1–3) were evaluated, along with an additional hybrid speciation scenario (4), which incorporated the intermediate position of *O. hikimius* between ncClade 1 and ncClade 2, as suggested by the Neighbour-net and admixture analyses. Among these, scenario 4 was best supported, suggesting that the two *O. obscurus* clades diverged first, followed by the emergence of *O. hikimius* through secondary contact between them (Fig. 5). Parameter estimation under scenario 4 indicated that the divergence of *O. obscurus* into two ncClades occurred approximately 2.18 Ma (95% HPD, 0.41–10.41 Ma) when assuming a generation time of three years, or 2.90 Ma (0.55–13.88) when assuming four years. The emergence of *O. hikimius* was estimated at approximately 1.63 Ma (0.36–5.65) or 2.17 Ma (0.48– 7.54). These SNP-based divergence time estimates were broadly consistent with those inferred from the mtDNA time tree, considering the overlapping confidence intervals, although they tended to suggest a more recent divergence event.

**Fig. 5.**
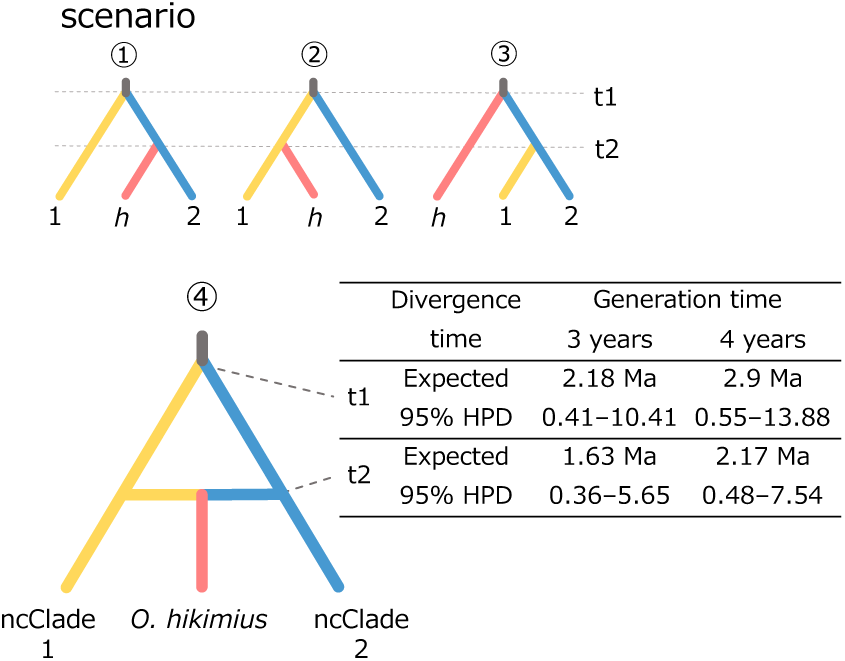
Four divergence scenarios among *Odontobutis hikimius* and the two ncClades of *O. obscurus*, with scenario 4 selected as the best supported. Estimated times of divergence and gene flow are shown for scenario 4

## 4. Discussion

### 4.1 Effective phylogeographic survey combining eDNA and genome-wide SNP analyses

The two-step survey strategy, combining eDNA analysis and genome-wide SNP analysis, effectively revealed that Japanese *O. obscurus* comprises deeply divergent lineages, comparable in divergence depth to its split from *O. hikimius*, each of which exhibits geographically fine-scale genetic population structures. This success stems from the complementary advantages of each method: the simplicity and broad applicability of eDNA surveys (Tsuji et al., 2023), and the detailed genetic resolution provided by PCR-based SNP analysis (Fujimoto et al., 2022). For the eDNA survey in this study, water sampling was conducted at a total of 338 sites, including 54 (16%) used in a previous study (Tsuji et al., 2023). All sampling was conducted through several short trips by one or two investigators or via water sampling and postal shipment by local collaborators. Conducting a capture-based survey on a comparable scale would be impractical due to the labour, time, and financial costs involved.

Dense eDNA sampling over a wide area is likely to maximise the detection of local genetic groups, particularly in species with high regional specificity such as *Odontobutis* species. For mtClade 2, which was previously regarded as a single local genetic group (Mukai and Nishida, 2003; Sakai et al., 1998), the surveys conducted across a large number of sites in this study revealed greater within-clade diversity and identified several distinct local population groups. Furthermore, the Tango group within this clade, newly identified in this study, had been overlooked in previous studies due to their limited spatial coverage, which did not encompass its distributional range. These findings indicate that eDNA surveys conducted across a large number of sites can incidentally detect previously unrecognised regional populations. In addition, the reusability of archived samples makes eDNA-based phylogeographic surveys increasingly cost-effective. All samples used in this study can be reused for future studies on different species, so additional sampling is expected to be minimal.

Although eDNA generally provides limited phylogeographic resolution due to the short sequence lengths available, genome-wide SNP analysis, employing strategic population sampling guided by eDNA results for capture surveys, enables the elucidation of finer phylogeographic patterns at minimal cost. Our PCR-based SNP analysis effectively elucidated the overall picture of population divergence and phylogenetic relationships in *Odontobutis* species in Japan. PCR-based SNP analysis, such as MAAS and MIG-seq, typically yields fewer SNPs (several hundred to a few thousand) than larger-scale reduced-representation sequencing methods, such as RAD-seq and GRAS-Di, which can generate tens to hundreds of thousands of SNPs. Nevertheless, the former generally provides sufficient phylogeographic resolution to detect genetic differences among regional populations. An additional major advantage of PCR-based SNP analysis is the simplicity and efficiency of the library preparation process, which requires only a small amount of tissue or DNA materials (Fujimoto et al., 2022; Suyama and Matsuki, 2015). In this study, MAAS library preparation of DNA extracted from 177 specimens was completed by a single researcher (S.T.) in practically one day (two afternoons). The likelihood of successfully capturing target species in the field can also be increased by selecting survey sites where high eDNA concentrations were detected in prior eDNA analyses, thereby reducing the labour, time and cost challenges associated with capture surveys.

The acquisition of longer DNA sequences from eDNA is one of the key future research priorities. Although our results, together with those of a few previous studies (Tsuji et al., 2023; Yatsuyanagi et al., 2024), demonstrate that short eDNA sequences (≤400 bp) are sufficiently powerful to outline the population structure of target species, longer sequence data are required for more detailed population genetic and phylogenetic analyses. To meet this need, this and previous studies additionally obtained longer sequences through tissue-based mtDNA sequencing, which remains labour-intensive and time-consuming, particularly given the large number of specimens typically required. Developing techniques to analyse longer sequences directly from eDNA samples will eliminate the reliance on tissue-based sequencing, thereby enabling more efficient and cost-effective phylogeographic surveys. Currently, eDNA analysis is optimised for detecting shorter sequences (≤400 bp); however, recent findings suggest that longer mtDNA fragments (≥1,000 bp) are also present in environmental samples (Doorenspleet et al., 2025; Jensen et al., 2021; Maggini et al., 2024; Wang et al., 2021). With recent advances in high-precision sequencing technologies, such as Nanopore sequencing, the analysis of longer DNA fragments from eDNA samples is likely to become feasible in the near future.

### 4.2 *Hybrid origin of* Odontobutis hikimius

*Odontobutis hikimius* was described relatively recently as a distinct species, separated from *O. obscurus* based on morphological characteristics (e.g., features of the cephalic sensory system) and genetic distinctiveness (Iwata and Sakai, 2002; Sakai et al., 1998). However, its origin has not been studied in detail. The selected divergence scenario in this study suggested that *O. hikimius* originated through hybridisation, following secondary contact between ncClade 1 and ncClade 2 of *O. obscurus*. Although the inferred phylogenetic relationships among *O. hikimius* and the two major clades of *O. obscurus* differed between the mtDNA and SNP trees, both indicated a near-trifurcation with low statistical support. Admixture analysis with K = 2–3 also supported the presence of genetic components from both major clades of *O. obscurus* in *O. hikimius*. Furthermore, the fact that the distribution range of *O. hikimius* is located at the boundary between the current distribution ranges of ncClade 1 and ncClade 2 provides additional support to the hybrid origin of *O. hikimius*. Although more detailed genomic investigation is still needed to confirm the hybrid origin of *O. hikimius*, the present results indicate a high likelihood that this species represents the first documented case of hybrid speciation among Japanese freshwater fishes. Intriguingly, a hybrid zone between *O. hikimius* and ncClade 1-3 of *O. obscurus* was identified in the middle reaches in the Takatsu River system. The gene flow observed in this restricted area suggests that reproductive isolation is sufficiently effective to maintain genetic separation between *O. hikimius* populations in the upper reaches and *O. obscurus*, although it may not be complete. Further research on the extent and mechanisms of their reproductive isolation is necessary.

The divergence time between the two major clades of *O. obscurus* was estimated at 2.06–5.90 Ma (mtDNA) or 0.41–13.88 Ma (demographic modelling based on SNPs, assuming the generation time of 3 or 4 years). Then, the secondary contact between the two clades was estimated to have occurred 1.59–4.76 Ma (mtDNA) and 0.36–7.54 Ma (SNPs). Due to the deep timescale of these events and the large confidence intervals, it is difficult to infer the geographical factors responsible for the divergence of the major clades and subsequent secondary contact. However, these events likely occurred during the early stages of freshwater fish fauna formation in Japan (Watanabe et al., 2017). The two major clades of *O. obscurus*, which are inferred to have given rise to *O. hikimius*, may have diverged at a species level, and further taxonomic investigation is therefore warranted.

### 4.3 Phylogeographic patterns of Odontobutis linked to geological history

The detailed phylogeographic patterns of *Odontobutis* revealed by the present eDNA and SNP analyses strongly support several previously reported, albeit fragmented, distributional patterns of freshwater fishes in the Japanese archipelago. Together with the few existing large-scale phylogeographic studies based on mtDNA (e.g., Nakagawa et al., 2016; Tominaga et al., 2016), they offer a comprehensive biogeographic basis for understanding the formation of freshwater biota in western Japan. The two clades of *O. obscurus* identified in this study were primarily distributed in the eastern and western regions, respectively. The haplotypes of two clades were rarely detected in the same site, and their distribution boundaries were clearly demarcated by river systems and mountain ranges. This suggests that past geographical barriers remain intact today, with little gene flow occurring between populations associated with the two clades.

Both ncClade 1 and ncClade 2 consisted of several geographically cohesive subgroups, the distribution boundaries of which can largely be accounted for by mountain ranges, ancient river systems and the historical river capture events. The Japanese archipelago has experienced active orogenesis and repeated land connections with the continent since its formation in the Early Miocene (Marsaglia et al., 1992; Maruyama et al., 1997; Yonekura, 2001). Such geological events have been recognised as key drivers forming phylogeographic patterns of Japanese freshwater fishes (Watanabe et al., 2017). In particular, the Chugoku and Ibuki–Suzuka Mountains have been suggested to have acted as geographical barriers during the range expansion for several freshwater fishes (Fig. 4C; Watanabe et al., 2017; Watanabe and Takahashi, 2010). In *O. obscurus* as well, these mountain ranges served as boundaries between regional genetic groups, with the Chugoku Mountains separating ncClade 2-7 and other ncClade 2 groups and the Ibuki– Suzuka Mountains dividing ncClades 2-4 and 2-5. Although the precise timing of the uplift of the Chugoku Mountains remains unclear, it has been estimated to have started in the Late Pliocene to Early Pleistocene (Fujita 1985; Kitamura 2010). This is consistent with our results, which suggest that ncClade2-7 (mtDNA from E. San’in and W. San’in) diverged from the other groups approximately 1.99 Ma (95% HPD, 0.99–3.10 Ma). Furthermore, the uplift of the Ibuki–Suzuka Mountains has been estimated to have started around 1.0–1.5 Ma (Kawabe, 1989), which is generally consistent with, but slightly later than, the estimated divergence time of Tokai (ncClade 2-4) and LakeBiwa (ncClade 2-5) groups (0.44 Ma; 0.19–0.72 Ma).

The distribution of regional groups on the northern and southern sides of the Seto Inland Sea suggests that palaeoriver systems, repeatedly formed in the present-day sea area during glacial low sea-level periods (Ota et al., 2004; Sato and Yasuda, 2022), would have served as dispersal corridors for *O. obscurus*. The Houyo and Kitan palaeoriver systems once flowed westward and eastward across the Seto Inland Sea area, with their northern and southern tributaries connecting river systems on Honshu and Shikoku Islands (Fig. 4C). The current disjunct distributions of ncClade 1-3, 1-4, 2-1, and 2-6, each found across the northern and southern sides of the Seto Inland Sea, are well explained by the courses of these palaeoriver systems. Similar distribution patterns have been reported in several other freshwater fishes (Takahashi et al., 2020; Watanabe et al., 2014), and our findings further emphasise the important role of palaeoriver systems in shaping the freshwater fish fauna and population structures in western Japan.

The phylogeographic patterns of *O. obscurus* also highlight the importance of river capture events in shaping freshwater fish distribution within the mountainous island landscape of the Japanese archipelago. ncClade 1-4 is predominantly distributed around the Seto Inland Sea; however, some populations (9_Maruyama and 2_Wakasa) are found beyond the watershed boundaries on the Sea of Japan side, extending into the distribution range of ncClade 2. The upper reaches of the Kako River system, which drains into the Seto Inland Sea, have a complex history of repeated connections with the Maruyama and Yura River systems, both draining into the Sea of Japan, and have altered the flow direction northward through river capture events (Fig. 1B; Okada and Takahashi, 1969). Therefore, the distribution of ncClade 1-4 across the watershed boundaries is inferred to have resulted from range expansion following such river course changes. This hypothesis is further supported by similar distribution patterns observed in other freshwater species, including the cyprinid *Tanakia limbata* and *Hemibarbus barbus*, the loach *Lefua echigonia* and *Cobitis striata striata* (see Mizuno, 1977), the goby *Rhinogobius tyoni* (see Kunimatsu and Fuke, 2021), and the shrimp *Neocaridina denticulata* (see Fujita et al., 2011). Collectively, these examples detected in the phylogeographic patterns of *O. obscurus* underscore the dynamic processes that have shaped regional freshwater faunas, driven by the active tectonics of the Japanese archipelago.

### 4.4 Genetic disturbance of O. obscurus

The unnatural distributions observed in some regional groups through eDNA analysis (W. Seto, W. San’in, Lake Biwa and Hokuriku groups) suggest that genetic disturbance of *O. obscurus* due to human-mediated introductions may be occurring sporadically. Although there is no definitive evidence that the co-occurrence and hybridisation between ncClade 1 and 2 individuals found at st. 6*_Kino_Hashimoto is anthropogenic, their presence indicates that *O. obscurus* individuals from different regions could hybridise when they occur sympatrically.

Artificial introductions of species and of genetically distinct groups from other regions are causing serious problems for the conservation of freshwater ecosystems worldwide (Cucherousset and Olden, 2011; Gozlan et al., 2010). Although the establishment of *O. obscurus* populations in non-native areas within Japan has been reported in several cases (Hosoya, 2019; Kuroda, 1960; Mukai and Nishida, 2003; Yamaguti, 1939), reliable data on introductions and genetic disturbances within its native range have been lacking. The sporadic occurrence of geographically distant genetic groups across both native and non-native areas observed in this study highlights the ongoing domestic spread of *O. obscurus* in Japan. Furthermore, the presence of various genetic groups in diverse locations suggests that the introductions are more likely attributable to unregulated releases by aquarium hobbyists and shops, rather than to a single systematic human action (e.g., unintentional contamination associated with fisheries releases), as also suggested by previous reports (Mukai and Nishida, 2003; Yagyu et al., 2021). As discussed above, even deeply divergent groups such as ncClades 1 and 2 of *O. obscurus* exhibit limited reproductive isolation when artificially brought into contact. To conserve native genetic populations and prevent further genetic disturbance, a comprehensive assessment of the current extent and impact of such disturbance is necessary.

## Authors’ contributions

S.T. and K.W. conceived and designed the research. S.T., Y.M. and N.S. performed a field survey.

S.T. performed all molecular experiments. S.T., Y.M. and N.S. performed data analysis. S.T. and K.W. wrote the early draft and completed it with significant input from all authors.

## Data Availability Statement

All 12S rRNA to the first half of 16S rRNA sequence data newly obtained from tissue DNA in this study are available from DDBJ under accession numbers: LC905600-LC905775 (Table S2). All raw sequences and metadata are deposited in the DDBJ Sequence Read Archive with the accession numbers: DRR794066–DRR794328 (eDNA data); DRR802370–DRR802546 (MAAS data).

## Supporting information

Supplementary_Further methodological details

## Acknowledgements

We thank our cooperators and laboratory members for providing eDNA and tissue samples: S. Kunimatsu, E. Agyeman, A. Kogayu (Kyoto University), Y. Fuke (Setsunan University), R. Inui (Fukuoka Institute of Technology), R. Nakao (Yamaguchi University), Y. Oto (Ehime University), S. Seki (Biwako Base). We thank K. Onuki, K. Ido (Kyoto University) and Y. Fuke for helping analysis of SNP data. This study was financially supported by ESPEC Foundation for Global Environment Research and Technology (Charitable Trust) and the Sasagawa Scientific Research Grant from the Japan Science Society.

## Figure legends

Fig. S1 The results of ADMIXTURE analysis using genome-wide 663 SNPs obtained from all specimens of Japanese *Odontobutis* species (n = 176): (A) Relationship between cross-validation error (CV-error) and the number of clusters (K), and (B) Ancestry bar plots for each K. The asterisks (*1 and *2) indicate the sites where hybrid individuals inhabit, 18_Takatsu_MD and 6_Kino_Hashimoto, respectively (see Figure 1C).

Fig. S2 The results of ADMIXTURE analysis using genome-wide 663 SNPs obtained from 74 specimens assigned to ncClade2 in Fig. 4: (A) Relationship between cross-validation error (CV-error) and the number of clusters (K), and (B) Ancestry bar plots for each K.

Fig. S3 The results of ADMIXTURE analysis using genome-wide 663 SNPs obtained from 89 specimens assigned to ncClade2 in Fig. 4: (A) Relationship between cross-validation error (CV-error) and the number of clusters (K), and (B) Ancestry bar plots for each K.

## References

Alexander, D.H., Novembre, J., Lange, K., 2009. Fast model-based estimation of ancestry in unrelated individuals. Genome Reserch 19, 1655–1664. 10.1101/gr.094052.109

Almeida-Rocha, J.M., Soares, L.A.S.S., Andrade, E.R., Gaiotto, F.A., Cazetta, E., 2020. The impact of anthropogenic disturbances on the genetic diversity of terrestrial species: A global meta-analysis. Molecular Ecology 29, 4812–4822. 10.1111/mec.15688

Antich, A., Palacín, C., Zarcero, J., Wangensteen, O.S., Turon, X., 2023. Metabarcoding reveals high-resolution biogeographical and metaphylogeographical patterns through marine barriers. Journal of Biogeography 50, 515–527. 10.1111/jbi.14548

Avise, J.C., 2000. Phylogeography: the history and formation of species. Harvard University Press, Cambridge.

Baird, N.A., Etter, P.D., Atwood, T.S., Currey, M.C., Shiver, A.L., Lewis, Z.A., Selker, E.U., Cresko, W.A., Johnson, E.A., 2008. Rapid SNP discovery and genetic mapping using sequenced RAD markers. PLOS ONE 3, e3376. 10.1371/journal.pone.0003376

Ballard, J.W.O., Whitlock, M.C., 2004. The incomplete natural history of mitochondria. Molecular Ecology 13, 729–744. 10.1046/j.1365-294X.2003.02063.x

Becker, R.A., Wilks, A.R., Brownrigg, R., Minka, T.P., Deckmyn, A., 2021. maps: Draw Geographical Maps. ver. 3.4.0.

Beheregaray, L.B., 2008. Twenty years of phylogeography: the state of the field and the challenges for the Southern Hemisphere. Molecular Ecology 17, 3754–3774. 10.1111/j.1365-294X.2008.03857.x

Bermingham, E., Moritz, C., 1998. Comparative phylogeography: concepts and applications. Molecular Ecology 7, 367–369. 10.1046/j.1365-294x.1998.00424.x

Bowen, B.W., 2016. The three domains of conservation genetics: Case histories from Hawaiian waters. Journal of Heredity 107, 309–317. 10.1093/jhered/esw018

Brownrigg, R., 2018. Extra Map Databases. Version 2.3.0 [R package].

Callahan, B.J., McMurdie, P.J., Rosen, M.J., Han, A.W., Johnson, A.J.A., Holmes, S.P., 2016. DADA2: High-resolution sample inference from Illumina amplicon data. Nature Methods 13, 581–583. 10.1038/nmeth.3869

Chen, S., Zhou, Y., Chen, Y., Gu, J., 2018. fastp: an ultra-fast all-in-one FASTQ preprocessor. Bioinformatics 34, i884–i890. 10.1093/bioinformatics/bty560

Clement, M., Posada, D., Crandall, K.A., 2000. TCS: a computer program to estimate gene genealogies. Molecular Ecology 9, 1657–1659. 10.1046/j.1365-294x.2000.01020.x

Collin, F.-D., Durif, G., Raynal, L., Lombaert, E., Gautier, M., Vitalis, R., Marin, J.-M., Estoup, A., 2021. Extending approximate Bayesian computation with supervised machine learning to infer demographic history from genetic polymorphisms using DIYABC Random Forest. Molecular Ecology Resources 21, 2598–2613. 10.1111/1755-0998.13413

Couton, M., Viard, F., Altermatt, F., 2023. Opportunities and inherent limits of using environmental DNA for population genetics. Environmental DNA 5, 1048–1064. 10.1002/edn3.448

Cucherousset, J., Olden, J.D., 2011. Ecological impacts of nonnative freshwater fishes. Fisheries 36, 215–230. 10.1080/03632415.2011.574578

Doorenspleet, K., Jansen, L., Oosterbroek, S., Kamermans, P., Bos, O., Wurz, E., Murk, A., Nijland, R., 2025. The long and the short of it: Nanopore-based eDNA metabarcoding of marine vertebrates works; sensitivity and species-level assignment depend on amplicon lengths. Molecular Ecology Resources 25, e14079. 10.1111/1755-0998.14079

Enoki, H., Takeuchi, Y., 2018. New genotyping technology, GRAS-Di, using next generation sequencer. Presented at the Plant and Animal Genome XXVI Conference (January 13 - 17, 2018), PAG.

Fujimoto, S., Yaguchi, H., Myosho, T., Aoyama, H., Sato, Y., Kimura, R., 2022. Population admixtures in medaka inferred by multiple arbitrary amplicon sequencing. Scientific Reports 12, 19989. 10.1038/s41598-022-24498-7

Fujita, J., Nakayama, K., Kai, Y., Ueno, M., Yamashita, Y., 2011. Geographical distributions of mitochondrial DNA lineages reflect ancient directions of river flow: A case study of the Japanese freshwater shrimp *Neocaridina denticulata denticulata* (Decapoda: Atyidae). jzoo 28, 712–718. 10.2108/zsj.28.712

Garrick, R.C., Bonatelli, I.A.S., Hyseni, C., Morales, A., Pelletier, T.A., Perez, M.F., Rice, E., Satler, J.D., Symula, R.E., Thomé, M.T.C., Carstens, B.C., 2015. The evolution of phylogeographic data sets. Molecular Ecology 24, 1164–1171. 10.1111/mec.13108

Gerritsen, H., 2018. Data Visualisation on Maps. ver. 1.5.1.

Gozlan, R.E., Britton, J.R., Cowx, I., Copp, G.H., 2010. Current knowledge on non-native freshwater fish introductions. Journal of Fish Biology 76, 751–786. 10.1111/j.1095-8649.2010.02566.x

Hamady, M., Walker, J.J., Harris, J.K., Gold, N.J., Knight, R., 2008. Error-correcting barcoded primers for pyrosequencing hundreds of samples in multiplex. Nature Methods 5, 235–237. 10.1038/nmeth.1184

Hickerson, M.J., Carstens, B.C., Cavender-Bares, J., Crandall, K.A., Graham, C.H., Johnson, J.B., Rissler, L., Victoriano, P.F., Yoder, A.D., 2010. Phylogeography’s past, present, and future: 10 years after Avise, 2000. Molecular Phylogenetics and Evolution 54, 291–301. 10.1016/j.ympev.2009.09.016

Hoang, D.T., Chernomor, O., Von Haeseler, A., Minh, B.Q., Vinh, L.S., 2018. UFBoot2: Improving the ultrafast bootstrap approximation. Molecular Biology and Evolution 35, 518–522. 10.1093/molbev/msx281

Hosoya, K., 2019. Yamakei Handy Illustrated Book 15: Freshwater fish of Japan, enlarged and revised edition. ed. Yama-kei Publishers co.,Ltd.

Hu, J., Li, H., Sakai, H., Mukai, T., Young Suk, H., Li, C., 2023. Molecular phylogenetics of the fresh water sleepers *Odontobutis* (Gobiiformes: Odontobutidae) and its implications on biogeography of freshwater ichthyofauna of East Asia. Molecular Phylogenetics and Evolution 186, 107871. 10.1016/j.ympev.2023.107871

Huson, D.H., Bryant, D., 2024. The SplitsTree App: interactive analysis and visualization using phylogenetic trees and networks. Nature Methods 21, 1773–1774. 10.1038/s41592-024-02406-3

Iwata, A., Sakai, H., 2002. *Odontobutis hikimius* n. sp.: A new freshwater goby from Japan, with a key to species of the genus. cope 2002, 104–110. 10.1643/0045-8511(2002)002%255B0104:OHNSAN%255D2.0.CO;2

Jense, C., Adams, M., Raadik, T.A., Waters, J.M., Morgan, D.L., Barmuta, L.A., Hardie, S.A., Deagle, B.E., Burridge, C.P., 2024. Cryptic diversity within two widespread diadromous freshwater fishes (Teleostei: Galaxiidae). Ecology and Evolution 14, e11201. 10.1002/ece3.11201

Jensen, M.R., Sigsgaard, E.E., Liu, S., Manica, A., Bach, S.S., Hansen, M.M., Møller, P.R., Thomsen, P.F., 2021. Genome-scale target capture of mitochondrial and nuclear environmental DNA from water samples. Molecular Ecology Resources 21, 690–702. 10.1111/1755-0998.13293

Kalkauskas, A., Perron, U., Sun, Y., Goldman, N., Baele, G., Guindon, S., Maio, N.D., 2021. Sampling bias and model choice in continuous phylogeography: Getting lost on a random walk. PLOS Computational Biology 17, e1008561. 10.1371/journal.pcbi.1008561

Kalyaanamoorthy, S., Minh, B.Q., Wong, T.K.F., von Haeseler, A., Jermiin, L.S., 2017. ModelFinder: fast model selection for accurate phylogenetic estimates. Nature Methods 14, 587–589. 10.1038/nmeth.4285

Katoh, K., Rozewicki, J., Yamada, K.D., 2019. MAFFT online service: multiple sequence alignment, interactive sequence choice and visualization. Briefings Bioinformatics 20, 1160–1166. 10.1093/bib/bbx108

Kawabe, T., 1989. Stratigraphy of the lower part of the Kobiwako group around the Ueno bosin, Kinki district, Japan. Journal of geosciences Osaka City University 32, 39–90.

Knowlton, N., 1993. Sibling species in the sea. Annual Review of Ecology and Systematics 24, 189–216.

Kunimatsu, S., Fuke, Y., 2021. First record of *Rhinogobius tyoni* (Gobiidae) from Yura River in Kyoto Prefecture, Japan and morphological comparison with *Rhinogobius* sp. OR. Ichthy, Natural History of Fishes of Japan 4, 12–17. 10.34583/ichthy.4.0_12

Kuroda, N., 1960. A new list of the fishes of Lake Suwa. Japanese Journal of Ichthyology 8, 35–46. 10.11369/jji1950.8.35

Leigh, J.W., Bryant, D., 2015. popart: full-feature software for haplotype network construction. Methods in Ecology and Evolution 6, 1110–1116. 10.1111/2041-210X.12410

Liu, X.P., Duffy, G.A., Pearman, W.S., Pertierra, L.R., Fraser, C.I., 2022. Meta-analysis of Antarctic phylogeography reveals strong sampling bias and critical knowledge gaps. Ecography 2022, e06312. 10.1111/ecog.06312

Maggini, S., Jacobsen, M.W., Urban, P., Hansen, B.K., Kielgast, J., Bekkevold, D., Jardim, E., Martinsohn, J.T., Carvalho, G.R., Nielsen, E.E., Papadopulos, A.S.T., 2024. Nanopore environmental DNA sequencing of catch water for estimating species composition in demersal bottom trawl fisheries. Environmental DNA 6, e555. 10.1002/edn3.555

Marsaglia, K.M., Ingersoll, R.V., Packer, B.M., 1992. Tectonic evolution of the Japanese islands as reflected in modal compositions of Cenozoic forearc and backarc sand and sandstone. Tectonics 11, 1028–1044. 10.1029/91TC03183

Maruyama, S., Isozaki, Y., Kimura, G., Terabayashi, M., 1997. Paleogeographic maps of the Japanese Islands: Plate tectonic synthesis from 750 Ma to the present. Island Arc 6, 121–142. 10.1111/j.1440-1738.1997.tb00043.x

Mashiko, K., Yamagishi, H., 1976. Postembryonal growth of an eleotrid goby, *Odontobutis obscurus* (Temminck et Schlegel), as related to the development of behaviour. Bulletin of the Japanese Society of Scientific Fisheries 42, 953–959.

Matsuda, T., 2024. Freshwater fishes of Fukui, 1st ed. Katsuki Bookstore.

Minh, B.Q., Schmidt, H.A., Chernomor, O., Schrempf, D., Woodhams, M.D., von Haeseler, A., Lanfear, R., 2020. IQ-TREE 2: New models and efficient methods for phylogenetic inference in the genomic era. Molecular Biology and Evolution 37, 1530–1534. 10.1093/molbev/msaa015

Mizuno, N., 1977. Report of a fish survey in Hikami District. Hikami 9, 8–104.

Moritz, C., 2002. Strategies to protect biological diversity and the evolutionary processes that sustain it. Systematic Biology 51, 238–254. 10.1080/10635150252899752

Mukai, T., Nishida, M., 2003. Mitochondrial DNA phylogeny of Japanese freshwater goby, *Odontobutis obscura*, and an evidence for artificial transplantation to Kanto District. Japanese Journal of Ichthyology 50, 71–76.

Nakagawa, H., Seki, S., Ishikawa, T., Watanabe, K., 2016. Genetic population structure of the Japanese torrent catfish *Liobagrus reinii* (Amblycipitidae) inferred from mitochondrial cytochrome *b* variations. Ichthyological Research 63, 333–346. 10.1007/s10228-015-0503-6

Nakajima, S., Tsuri, K., 2024. Testing the applicability of environmental DNA metabarcoding to landscape genetics. Molecular Ecology Resources 24, e13990. 10.1111/1755-0998.13990

Okada, A., Takahashi, K., 1969. Geomorphic development of the drainage basin of the river Yura, Western Honshu, Japan. Journal of Geography (Chigaku Zasshi) 78, 19–37. 10.5026/jgeography.78.19

Ota, Y., Naruse, T., Tanaka, S., Okada, A., 2004. Regional geomorphology of the Japanese Islands vol. 6 Geomorphology of Kinki Region. University of Tokyo Press.

Pedersen, M.W., De Sanctis, B., Saremi, N.F., Sikora, M., Puckett, E.E., Gu, Z., Moon, K.L., Kapp, J.D., Vinner, L., Vardanyan, Z., Ardelean, C.F., Arroyo-Cabrales, J., Cahill, J.A., Heintzman, P.D., Zazula, G., MacPhee, R.D.E., Shapiro, B., Durbin, R., Willerslev, E., 2021. Environmental genomics of Late Pleistocene black bears and giant short-faced bears. Current Biology 31, 2728–2736.e8. 10.1016/j.cub.2021.04.027

Pelletier, T.A., Parsons, D.J., Decker, S.K., Crouch, S., Franz, E., Ohrstrom, J., Carstens, B.C., 2022. phylogatR: Phylogeographic data aggregation and repurposing. Molecular Ecology Resources 22, 2830–2842. 10.1111/1755-0998.13673

Penn, O., Privman, E., Ashkenazy, H., Landan, G., Graur, D., Pupko, T., 2010. GUIDANCE: a web server for assessing alignment confidence scores. Nucleic Acids Res 38, W23–W28. 10.1093/nar/gkq443

Purcell, S., Neale, B., Todd-Brown, K., Thomas, L., Ferreira, M.A.R., Bender, D., Maller, J., Sklar, P., Bakker, P.I.W. de, Daly, M.J., Sham, P.C., 2007. PLINK: A tool set for whole-genome association and population-based linkage analyses. The American Journal of Human Genetics 81, 559–575. 10.1086/519795

R Core Team, 2023. R: A language and environment for statistical computing. R Foundation for Statistical Computing.

Rochette, N.C., Rivera-Colón, A.G., Catchen, J.M., 2019. Stacks 2: Analytical methods for paired-end sequencing improve RADseq-based population genomics. Molecular Ecology 28, 4737–4754. 10.1111/mec.15253

Rourke, M.L., Fowler, A.M., Hughes, J.M., Broadhurst, M.K., DiBattista, J.D., Fielder, S., Wilkes Walburn, J., Furlan, E.M., 2022. Environmental DNA (eDNA) as a tool for assessing fish biomass: A review of approaches and future considerations for resource surveys. Environmental DNA 4, 9–33. 10.1002/edn3.185

Rozas, J., Ferrer-Mata, A., Sánchez-DelBarrio, J.C., Guirao-Rico, S., Librado, P., Ramos-Onsins, S.E., Sánchez-Gracia, A., 2017. DnaSP 6: DNA sequence polymorphism analysis of large data sets. Molecular Biology and Evolution 34, 3299–3302. 10.1093/molbev/msx248

Sakai, H., Yamamoto, C., Iwata, A., 1998. Genetic divergence, variation and zoogeography of a freshwater goby, *Odontobutis obscura*. Ichthyological Research 45, 363–376. 10.1007/BF02725189

Sato, J.J., Yasuda, K., 2022. Ancient rivers shaped the current genetic diversity of the wood mouse (*Apodemus speciosus*) on the islands of the Seto Inland Sea, Japan. Zoological Letter 8, 9. 10.1186/s40851-022-00193-3

Scoble, J., Lowe, A.J., 2010. A case for incorporating phylogeography and landscape genetics into species distribution modelling approaches to improve climate adaptation and conservation planning. Diversity and Distributions 16, 343–353. 10.1111/j.1472-4642.2010.00658.x

Seebens, H., Blackburn, T.M., Dyer, E.E., Genovesi, P., Hulme, P.E., Jeschke, J.M., Pagad, S., Pyšek, P., van Kleunen, M., Winter, M., Ansong, M., Arianoutsou, M., Bacher, S., Blasius, B., Brockerhoff, E.G., Brundu, G., Capinha, C., Causton, C.E., Celesti-Grapow, L., Dawson, W., Dullinger, S., Economo, E.P., Fuentes, N., Guénard, B., Jäger, H., Kartesz, J., Kenis, M., Kühn, I., Lenzner, B., Liebhold, A.M., Mosena, A., Moser, D., Nentwig, W., Nishino, M., Pearman, D., Pergl, J., Rabitsch, W., Rojas-Sandoval, J., Roques, A., Rorke, S., Rossinelli, S., Roy, H.E., Scalera, R., Schindler, S., Štajerová, K., Tokarska-Guzik, B., Walker, K., Ward, D.F., Yamanaka, T., Essl, F., 2018. Global rise in emerging alien species results from increased accessibility of new source pools. Proceedings of the National Academy of Sciences 115, E2264–E2273. 10.1073/pnas.1719429115

Sersics, A.N., Cosacov, A., Cocucci, A.C., Johnson, L.A., Pozner, R., Avila, L.J., Sites, J.W., Jr., Morando, M., 2011. Emerging phylogeographical patterns of plants and terrestrial vertebrates from Patagonia. Biological Journal of the Linnean Society 103, 475–494. 10.1111/j.1095-8312.2011.01656.x

Suyama, Y., Matsuki, Y., 2015. MIG-seq: an effective PCR-based method for genome-wide single-nucleotide polymorphism genotyping using the next-generation sequencing platform. Scientific Reports 5, 16963. 10.1038/srep16963

Takahashi, T., Nagano, A.J., Kawaguchi, L., Onikura, N., Nakajima, J., Miyake, T., Suzuki, N., Kanoh, Y., Tsuruta, T., Tanimoto, T., Yasui, Y., Oshima, N., Kawamura, K., 2020. A ddRAD-based population genetics and phylogenetics of an endangered freshwater fish from Japan. Conservation Genetics 21, 641–652. 10.1007/s10592-020-01275-5

Templeton, A.R., Sing, C.F., 1993. A cladistic analysis of phenotypic associations with haplotypes inferred from restriction endonuclease mapping. IV. Nested analyses with cladogram uncertainty and recombination. Genetics 134, 659–669. 10.1093/genetics/134.2.659

Tsuji, S., Doi, H., Hibino, Y., Shibata, N., Watanabe, K., 2024. Rapid assessment of invasion front and biological impact of the invasive fish *Coreoperca herzi* using quantitative eDNA metabarcoding. Biological Invasions 26, 3107–3123. 10.1007/s10530-024-03364-9

Tsuji, S., Kunimatsu, S., Watanabe, K., 2025. Environmental DNA comparative phylogeography: simultaneous estimation of population structures within a species-rich group of freshwater gobies. Molecular Ecology Early view, e70059. 10.1111/mec.70059

Tsuji, S., Shibata, N., Inui, R., Nakao, R., Akamatsu, Y., Watanabe, K., 2023. Environmental DNA phylogeography: Successful reconstruction of phylogeographic patterns of multiple fish species from cups of water. Molecular Ecology Resources 23, 1050–1065. 10.1111/1755-0998.13772

Turchetto-Zolet, A.C., Pinheiro, F., Salgueiro, F., Palma-Silva, C., 2013. Phylogeographical patterns shed light on evolutionary process in South America. Molecular Ecology 22, 1193–1213. 10.1111/mec.12164

Turon, X., Antich, A., Palacín, C., Præbel, K., Wangensteen, O.S., 2020. From metabarcoding to metaphylogeography: separating the wheat from the chaff. Ecological Applications 30, e02036. 10.1002/eap.2036

Unmack, P.J., Cook, B.D., Johnson, J.B., Hammer, M.P., Adams, M., 2023. Phylogeography of a widespread Australian freshwater fish, western carp gudgeon (Eleotridae: *Hypseleotris klunzingeri*): Cryptic species, hybrid zones, and strong intra-specific divergences. Ecology and Evolution 13, e10682. 10.1002/ece3.10682

Wang, S., Yan, Z., Hänfling, B., Zheng, X., Wang, P., Fan, J., Li, J., 2021. Methodology of fish eDNA and its applications in ecology and environment. Science of The Total Environment 755, 142622. 10.1016/j.scitotenv.2020.142622

Watanabe, K., 2012. Faunal structure of Japanese freshwater fishes and its artificial disturbance. Environ Biol Fish 94, 533–547. 10.1007/s10641-010-9601-5

Watanabe, K., Mori, S., Tanaka, T., Kanagawa, N., Itai, T., Kitamura, J., Suzuki, N., Tominaga, K., Kakioka, R., Tabata, R., Abe, T., Tashiro, Y., Hashimoto, Y., Nakajima, J., Onikura, N., 2014. Genetic population structure of *Hemigrammocypris rasborella* (Cyprinidae) inferred from mtDNA sequences. Ichthyological Research 61, 352–360. 10.1007/s10228-014-0406-y

Watanabe, K., Takahashi, H., 2010. Natural history of freshwater fish geography. Hokkaido Unversity Press.

Watanabe, K., Tominaga, K., Nakajima, J., Kakioka, R., Tabata, R., 2017. Japanese Freshwater Fishes: Biogeography and Cryptic Diversity, in: Motokawa, M., Kajihara, H. (Eds.), Species diversity of animals in Japan. Springer Japan, Tokyo, pp. 183–227. 10.1007/978-4-431-56432-4_7

Yagyu, M., Nakamura, A., Mima, J., Oohara, H., 2021. The invasive Japanese freshwater goby *Odontobutis obscura* found in the tributary of Tenryu River Nagano prefecture, Japan. Natural History Reports of Inadani 22, 49–54. 10.20807/icmnhr.22.0_49

Yamaguti, S., 1939. Studies on the helminth fauna of Japan. Part 28. *Nipponaenia Chaenogobii*, a new cestode representing a new order from freshwater fishes. Japanese Journal of Zoology 8, 285–289.

Yamanaka, H., Minamoto, T., Matsuura, J., Sakurai, S., Tsuji, S., Motozawa, H., Hongo, M., Sogo, Y., Kakimi, N., Teramura, I., Sugita, M., Baba, M., Kondo, A., 2017. A simple method for preserving environmental DNA in water samples at ambient temperature by addition of cationic surfactant. Limnology 18, 233–241. 10.1007/s10201-016-0508-5

Yatsuyanagi, T., Kanbe, T., Fujii, K., Inoue, S., Araki, H., 2024. Environmental DNA unveils deep phylogeographic structure of a freshwater fish. Molecular Ecology 33. 10.1111/mec.17337

Yonekura, N., 2001. Regional geomorphology of the Japanese Islands, vol 1. Introduction to Japanese geomorphology. University of Tokyo Press, Tokyo.

