## Supplementary_Further methodological details for "High-resolution insights into the geographic differentiation and hybridisation of *Odontobutis* gobies through integrated eDNA and SNP analyses"

**Title:**

**Running title:**

Phylogeography of goby via eDNA and SNPs

**Authors:**

Satsuki Tsuji^1^*, Yugo Miuchi^2^, Naoki Shibata^3^, Katsutoshi Watanabe^2^ (*corresponding author)

**Affiliations:**

^1^ Biosphere Informatics Laboratory, Graduate School of Informatics, Kyoto University, Yoshidahonmachi, Sakyo-ku, Kyoto 606-8501, Japan

^2^ Graduate School of Science, Kyoto University, Kitashirakawa-Oiwakecho, Sakyo-ku, Kyoto 606-8502, Japan

^3^ Takara Bio Inc. 7-4-38 Nojihigashi, Kusatsu, Shiga 525-0058, Japan

**Corresponding author:**

Satsuki Tsuji

**Supplementary: Further methodological details**

***Environmental DNA extraction from filter samples***

DNA was extracted from each filter sample using a spin column method (EconoSpin, EP-31201; GeneDesign, Inc., Osaka, Japan) with the DNeasy Blood and Tissue Kit (Qiagen), following the protocol by Tsuji et al. (2024). The silica membrane was first removed from the spin column. All reagents, except for ethanol, were used as provided with the kit. Each filter was folded and placed in the spin column, which was then centrifuged at 5,000 g for 1 min to remove residual water, including benzalkonium chloride (Tsuji et al., 2022). Next, 475 μL of lysis buffer (200 μL ultrapure water, 250 μL Buffer AL, and 25 μL proteinase K) was added to the filter and incubated at 56°C for 40 min. After incubation, the column was centrifuged at 6,000 g for 1 min, and 500 μL of 99.9% ethanol was added to the filtrate. The DNA was purified using a DNeasy mini spin column according to the manufacturer’s instructions and finally eluted with 100 μL Buffer AE. The eDNA samples were stored at −20°C for subsequent analysis.

***Paired-end library preparation for eDNA analysis***

To avoid contamination, the laboratory was completely separated before and after the PCR step, and the experimenter was restricted from returning to the pre-PCR room on the same day. In this study, eDNA library preparation was performed according to the method in Tsuji et al. (2023). The first PCR (1st PCR) was performed to specifically amplify 366 bp of mtDNA 12S ribosomal RNA (rRNA) of *Odontobutis obscurus*, *O. hikimius* and three types of standard DNAs (Table S6) using group-specific primers (Odon_12S) with adapter sequences developed by Tsuji et al. (2023). The 1st PCR was performed in a 12-µL total volume of reaction mixture containing 6 µL of 2 × KAPA HiFi HotStart ReadyMix (KAPA Biosystems, MA, USA), 0.72 µL of each Odon_12S primer (10 µM), 2.56 µL of sterilised distilled H2O,1.0 μL standard DNA mix (consists of std1, 5 copies; std2, 25 copies; std3, 50 copies) and 1.0 µL eDNA template, with four replicates prepared for each eDNA sample. The sequences of Odon_12S primer combined sequencing primers (italic) and six random hexamers (N) are as follows: Odon_12 S primer-F (5′-*ACA CTC TTT CCC TAC ACG ACG CTC TTC CGA TCT* NNN NNN TAT ACG AGA GGC TCA AGC TGA T-3′), Odon_12 S primer-R (5′-*GTG ACT GGA GTT CAG ACG TGT GCT CTT CCG ATC T*NN NNN NGT TTT ACC AGT TTT GCT TAC TAT GG-3′). In all first PCR runs, four replicates of no-template control using ultrapure water instead of the template were prepared and treated as PCR negative controls. The thermal conditions for the first PCR were 5 min at 95°C, 45 cycles of 20 s at 98°C, 20 s at 60°C, 40 s at 72°C and 5 min at 72°C. The pooled 1st PCR products were purified using Sera-Mag SpeedBeads Carboxylate-Modified magnetic particles (Hydrophobic) (Cytiva) (× 0.8 beads). The purified first PCR products for each group were adjusted to 0.5 ng/μL and mixed in equal quantities (hereafter “first PCR product mix”) to use as the second-round PCR template (second PCR).

The second PCR was performed in a 12-μL total volume of reaction mixture containing 6.0 μL of 2 × KAPA HiFi HotStart ReadyMix, 2.0 μL of each primer with index (1.8 μM) and 2.0 μL of the purified first PCR product mix. The sequences of the second PCR primers with adapter sequences, eight indexes (X; Hamady et al., 2008) and sequencing primers (italic) were as follows: forward primer (5′-AAT GAT ACG GCG ACC ACC GAG ATC TAC AXX XXX XXX *ACA CTC TTT CCC TAC ACG ACG CTC TTC CCA TCT*-3′); reverse primer (5′-CAA GCA GAA GAC GGC ATA CGA GAT XXX XXX XX*G TGA CTG GAG TTC AGA CGT GTG CTC TTC CGA TCT*-3′). Unique indexes were applied to each sample to distinguish them from one another. The thermal conditions for the second PCR were 3 min at 95°C, 12 cycles of 20 s at 98°C, 15 s at 72°C and 5 min at 72°C. Following the second PCR, the products were pooled, and target bands were excised and purified using a 2% E-Gel SizeSelect agarose gel (Thermo Fisher Scientific, Massachusetts, USA). The final library was shipped to Novogene Co., Ltd. and sequenced on an Illumina NovaSeq 6000 using their gigabyte data purchase service (250 PE sequences).

***Bioinformatics processing and denoising for eDNA sequence data***

In this study, bioinformatics analysis was conducted using the methods outlined by Tsuji et al. (2023). The raw fastq file was denoised using the Divisive Amplicon Denoising Algorithm 2 (DADA2) package (v. 1.22; Callahan et al., 2016). First, primer sequences, random hexamers, low-quality, and unexpectedly short reads were removed. The error model was then trained to identify and correct indel mutations and substitutions in the dereplicated sequences. The paired reads were subsequently merged, and an amplicon sequence variant (ASV)–sample matrix was constructed. To obtain a standard line specific to each sample, linear regression analysis was conducted by plotting the read counts of internal standard DNAs against their known copy numbers (with the intercept fixed at zero). For each sample, the eDNA copy number of each ASV was converted using the sample-specific standard line: eDNA copy number = read count/regression slope of the sample-specific standard line (Table S7). For species assignment of the unique merged sequences, a local BLASTN search was performed using a reference database developed through the makeblastdb function (Camacho et al., 2009). The reference database included all sequences of *O. obscurus* and *O. hikimius* downloaded from the National Centre for Biotechnology Information (NCBI) database (https://www-ncbi-nlm-nih-gov.kyoto-u.idm.oclc.org/) and three standard DNAs. Sequences with a ≥96% identity were assigned to a species based on the top BLAST hits. Only these assigned sequences were used in the further denoising step.

To minimise false positives, a two-step data screening process was applied following the approach of Tsuji et al. (2023). First, sequences with a frequency of less than 1% within each sample were excluded. Next, sequences with a frequency less than half that of the most predominant sequence in each sample were also removed. Although this screening process may result in false negatives for minor haplotypes, it is expected to have only a minimal effect on the identification of major genetic lineages. The remaining sequences were treated as haplotypes present in each sample. Phylogenetic analyses were then conducted using these haplotype sequences, along with those from the reference database.

***Sequencing of longer mtDNA region from tissue DNA***

For high-throughput amplicon sequencing, the 1st PCR was performed using three primer sets designed in this study to amplify longer regions from the 12S rRNA to the first half of 16S rRNA region of *O. obscurus* and *O. hikimius*. The sequences of each primer are as follows:

Odon1216S_1-F (5′-ACA CTC TTT CCC TAC ACG ACG CTC TTC CGA TCT NNN NNN TAT ACG AGA GGC TCA AGC TGA T-3′); Odon1216S_1-R (5′-GTG ACT GGA GTT CAG ACG TGT GCT CTT CCG ATC TNN NNN NGT TTT ACC AGT TTT GCT TAC TAT GG-3′); Odon1216S_2-F (5′-ACA CTC TTT CCC TAC ACG ACG CTC TTC CGA TCT NNN NNN GTC AGC TTA CCC TAT AAG GGC C-3′), Odon1216S_2-R (5′-GTG ACT GGA GTT CAG ACG TGT GCT CTT CCG ATC TNN NNN NTT AGC TAC GCT CTG GTT TTT CCA AG-3′); Odon1216S_3-F (5′-ACA CTC TTT CCC TAC ACG ACG CTC TTC CGA TCT NNN NNN GGT AAG TGT ACC GGA AGG TGT A-3′), Odon1216S_3-R (5′-GTG ACT GGA GTT CAG ACG TGT GCT CTTC CGA TCT NNN NNN CCT TAT AAC TGG CTG CCT TGA AGT A-3′). These primer sets were designed such that the sequences of the three amplicons could be joined to form 1,031 bp of single continuous sequence (Odon1216S_1, 373 bp; Odon1216S_2, 294 bp; Odon1216S_33, 364 bp). In the 1st PCR, DNA amplification for each individual was performed separately using the primer sets Odon1216S_1 and _3 and the Odon1216S_2. The KAPA HiFi HotStart ReadyMix was employed as a total of 12 µL PCR reaction mixture containing 0.5 µM of each primer and 1 µL of tissue DNA. The thermal conditions were as follows: 3 min at 95°C, 20 cycles of 20 s at 98°C, 20 s at 60°C and 40 s at 72°C, and 5 min at 72°C. The 1st PCR products were diluted 100-fold in TE buffer (pH 8.0) and mixed in a 2 (amplicon of primer set 1 and 3):1 (amplicon of Odon1216S_2) ratio per individual (defined as amplicon mix). In the 2nd PCR, index primers with adapter sequences were used to add a unique, individual-specific index sequence to the two products derived from the same individual, amplified in different PCR reactions (Hamady et al., 2008). The KAPA HiFi HotStart ReadyMix was employed as a total of 12 µL PCR reaction mixture containing 0.3 µM of each index primer and 1 µL of amplicon mix. The thermal conditions were as follows: 3 min at 95°C, 10 cycles of 30 s at 98°C and 30 s at 72°C, and 5 min at 72°C. Following the 2nd PCR, the products were pooled and purified using Sera-Mag SpeedBeads. The final library was shipped to Novogene Co., Ltd. and sequenced on an Illumina NovaSeq 6000 using their gigabyte data purchase service (250 PE sequences).

For Sanger sequencing, 1,090 bp of 12S rRNA to 16S rRNA that completely encompassed the sequence obtained by high-throughput amplicon sequencing were amplified using primers developed in this study. The primer sequence is as follows: Odon1216_Sanger-F (5′-GTA AAA CTC GTG CCA GCC AC-3′), Odon1216_Sanger-R (5′-CAC TCT TTT GCC ACA GAG ACG-3′). The TaKaRa Ex Taq HS (Takara, Shiga, Japan) was employed as a total of 20 µL PCR reaction mixture containing 2 µL of 10× Ex Taq buffer, 1.6 µL of dNTP mixture, 0.5 µM of each primer and 1 µL tissue DNA. The thermal conditions were as follows: 2 min at 94°C and 30 cycles of 30 s at 94°C, 25 s at 58°C and 60 s at 72°C, and 5 min at 72°C. Then, PCR amplicons were purified using Sera-Mag SpeedBeads. Sanger sequencing was performed by a commercial Sanger-sequencing service (Macrogen Japan Corp., Tokyo, Japan).

***Bioinformatics processing for longer mtDNA data obtained from tissue DNA***

The fastq file was quality-filtered using fastp v0.24.0 (Chen et al., 2018) with a quality threshold of >Q30, polyG trimming, and other parameters set to default. The primer sequences were removed using cutadapt v4.9, with the mismatch tolerance option (-e) set to 0.1, and sequences less than 220 bp were discarded. Sequence alignments were performed for each specimen using four mapping tools, minimap2, bwa-mem2, bowtie2, and snap-aligner-against, the complete mitochondrial genome of *Odontobutis obscura* (DDBJ accession no. MW646297) as the reference. Consensus sequences were obtained from the bam files generated by each mapping tool using the samtools consensus. The four consensus sequences obtained for each individual were aligned using the mafft (Katoh et al., 2019). If different bases were called among four mapping tools, the base called by the most tools was adopted and one sequence per specimen was determined. Finally, the resulting sequences were aligned with the reference sequence, and 111 consecutive bases that could not be consistently determined across specimens were removed.

**References**

Callahan, B. J., McMurdie, P. J., Rosen, M. J., Han, A. W., Johnson, A. J. A., & Holmes, S. P. (2016). DADA2: High-resolution sample inference from Illumina amplicon data. Nature Methods, 13(7), Article 7. https://doi.org/10.1038/nmeth.3869

Camacho, C., Coulouris, G., Avagyan, V., Ma, N., Papadopoulos, J., Bealer, K., & Madden, T. L. (2009). BLAST+: Architecture and applications. BMC Bioinformatics, 10(1), 421. https://doi.org/10.1186/1471-2105-10-421

Hamady, M., Walker, J. J., Harris, J. K., Gold, N. J., & Knight, R. (2008). Error-correcting barcoded primers for pyrosequencing hundreds of samples in multiplex. Nature Methods, 5(3), Article 3. https://doi.org/10.1038/nmeth.1184

Katoh, K., Rozewicki, J., & Yamada, K. D. (2019). MAFFT online service: Multiple sequence alignment, interactive sequence choice and visualization. Briefings in Bioinformatics, 20(4), 1160–1166. https://doi.org/10.1093/bib/bbx108

Tsuji, S., Doi, H., Hibino, Y., Shibata, N., & Watanabe, K. (2024). Rapid assessment of invasion front and biological impact of the invasive fish *Coreoperca herzi* using quantitative eDNA metabarcoding. Biological Invasions, 26(9), 3107–3123. https://doi.org/10.1007/s10530-024-03364-9

Tsuji, S., Nakao, R., Saito, M., Minamoto, T., & Akamatsu, Y. (2022). Pre-centrifugation before DNA extraction mitigates extraction efficiency reduction of environmental DNA caused by the preservative solution (benzalkonium chloride) remaining in the filters. Limnology, 23(1), 9–16. https://doi.org/10.1007/s10201-021-00676-w

Tsuji, S., Shibata, N., Inui, R., Nakao, R., Akamatsu, Y., & Watanabe, K. (2023). Environmental DNA phylogeography: Successful reconstruction of phylogeographic patterns of multiple fish species from cups of water. Molecular Ecology Resources, 23(5), 1050–1065. https://doi.org/10.1111/1755-0998.13772
