## Supplementary figures and images for "High-resolution insights into the geographic differentiation and hybridisation of *Odontobutis* gobies through integrated eDNA and SNP analyses"

### Supplemental Figures

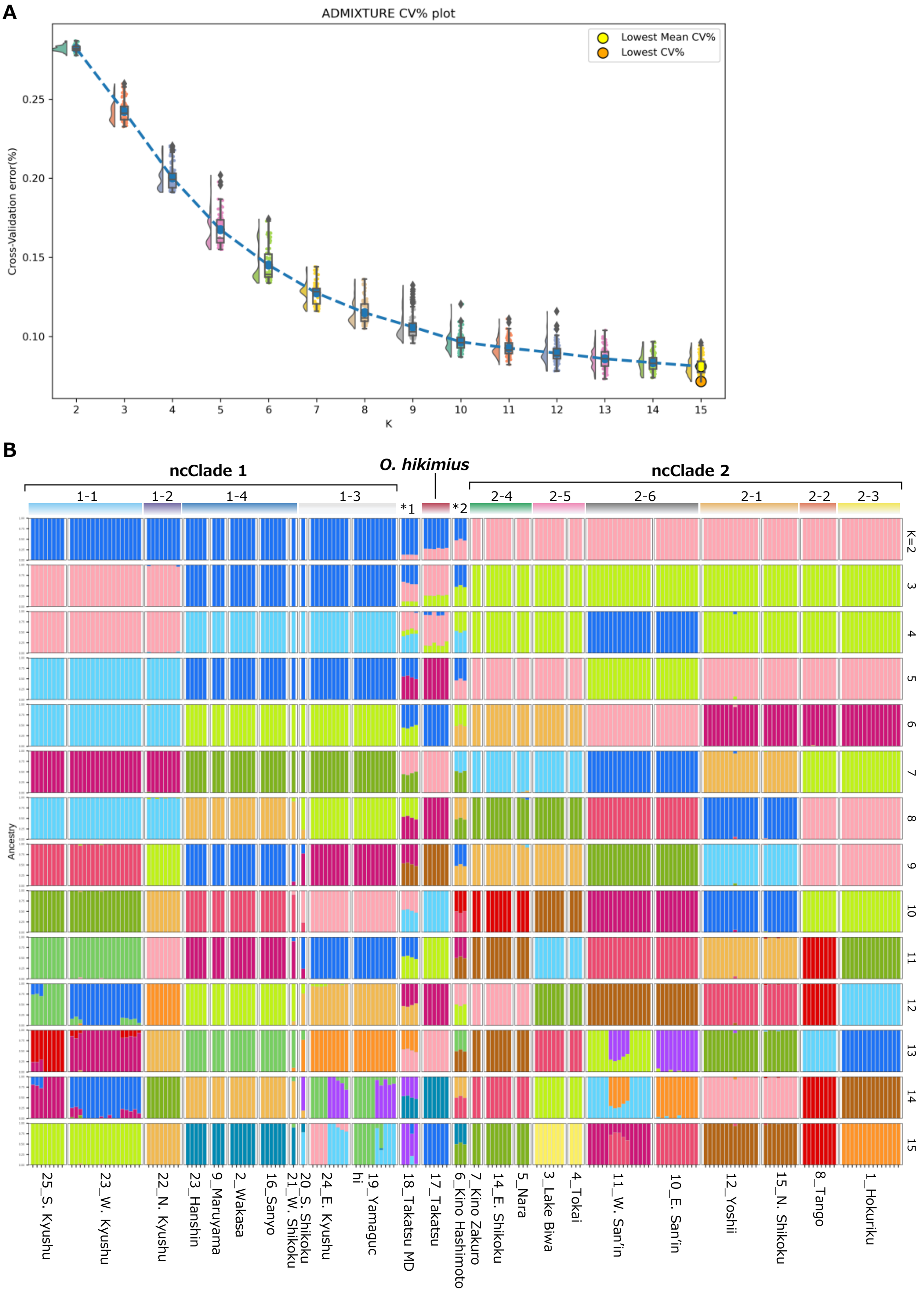

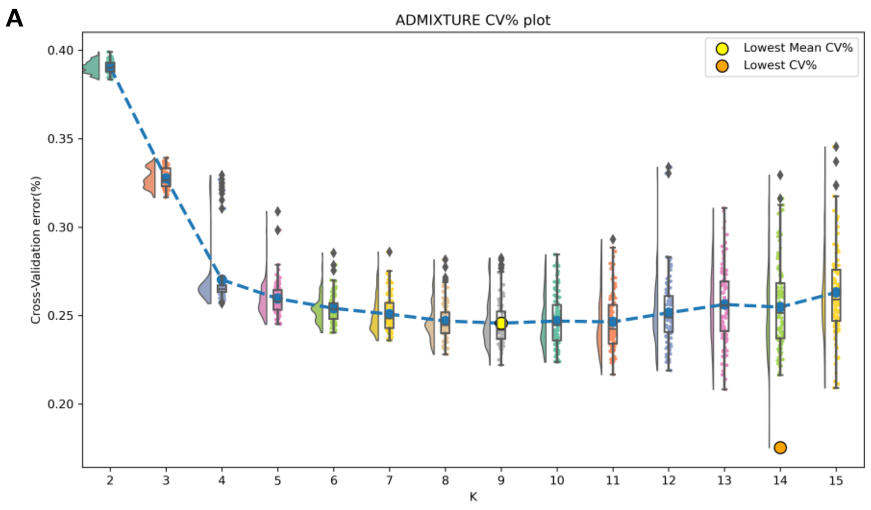

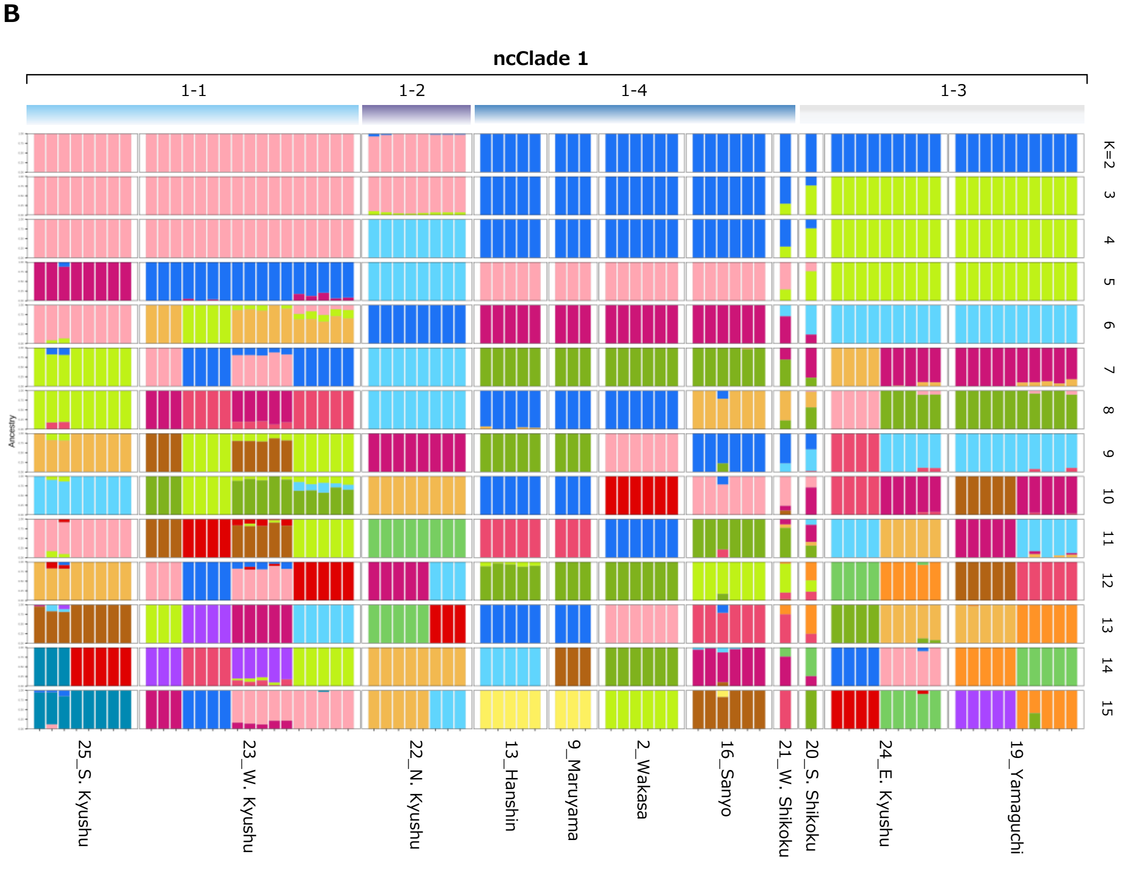

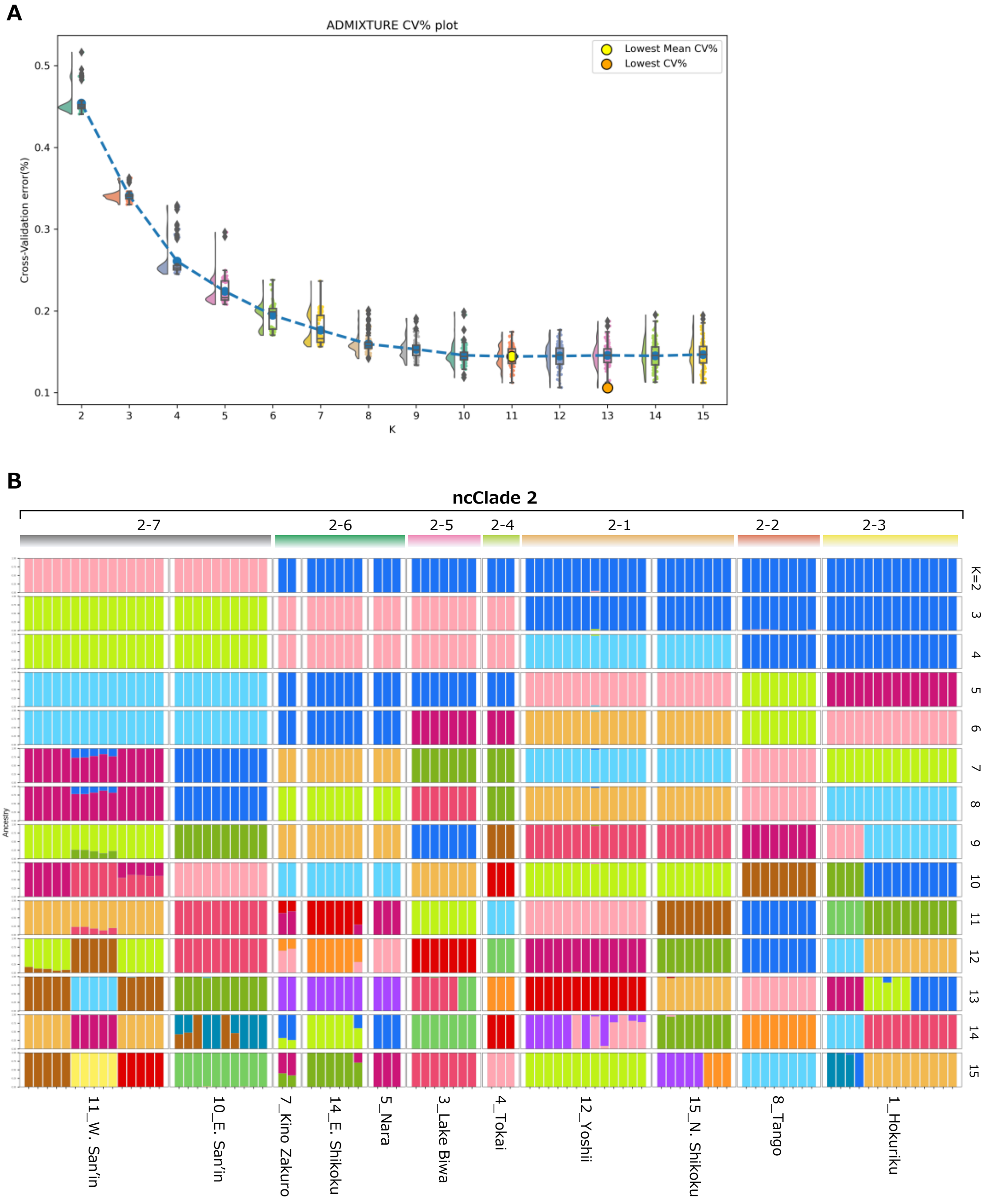
